# Drug-bound P-glycoprotein reveals nucleotide-dependent ligand pockets and ingress channels

**DOI:** 10.64898/2025.12.04.692475

**Authors:** Sungho Bosco Han, Jim Warwicker, Stephen M. Prince, Hao Fan

## Abstract

P-glycoprotein, an ATP-binding cassette exporter, coordinates drug recognition in a central cavity with nucleotide handling at nucleotide binding domains, yet the sequence and coupling of these events remain incompletely defined. Here, we report an integrated computational analysis of inward facing P-glycoprotein ensembles across varying drug locations, nucleotide states, and starting conformations. The sampled inward-facing ensembles of P-glycoprotein display continuous ligand cavity, reshaped by nucleotide occupancy, which enables polyspecific ligand recognition while transmitting allosteric signals to the nucleotide binding domains through intracellular coupling loops. Asymmetry between the two nucleotide binding sites follows from this communication, which alters the nucleotide coordination to potentially affect the catalytic progression. Our simulations reveal dynamic cavity access routes from cytosol and inner membrane, together with transient secondary structure within the interdomain flexible linker. Overall, our study gives insights into how ligand redistribution, asymmetric nucleotide coordination, and linker flexibility may correlate to form a unified view of early transport steps, yielding hypotheses for mutational and kinetic evaluation and informing strategies for small-molecule modulation of transporter function in cellular contexts where multidrug transport affects therapeutic response.

## 1 Introduction

P-glycoprotein (P-gp, ABCB1/MDR1) is a ubiquitous ATP-binding cassette (ABC) efflux transporter that protects cells by exporting a broad spectrum of xenobiotic compounds [1, 2, 3, 4]. This polyspecificity underlies the role of P-gp in multi-drug resistance, where it can recognize and extrude diverse chemotherapeutics ranging from small amphipathic drugs to bulky natural products such as the calcium channel blocker verapamil and the anti-mitotic alkaloid vinblastine [5, 6]. Notably, the distinction between “substrates” (transported compounds) and “inhibitors” of P-gp is often blurred: some classical P-gp inhibitors are actually transported at low concentrations, while many transportable substrates inhibit the ATPase activity of P-gp at higher concentrations [7, 8]. For instance, verapamil can both stimulate ATP hydrolysis and, at higher levels, suppress the transporter activity[9, 10]. Such biphasic drug–transporter interactions hits at the existence of multiple ligand binding sites and polyspecificity within the drug-binding pocket of P-gp. These features complicate efforts to categorize ligands strictly as substrates or inhibitors and underscore the need to understand how P-gp accommodates multiple drugs and modulates the catalytic cycle of P-gp in response. Understanding the mechanistic basis of the substrate polyspecificity of P-gp and drug-dependent ATPase modulation is not only fundamental to transporter biology, but also vital for improving pharmacotherapy and overcoming P-gp–mediated multi-drug resistance in disease.

Structurally, P-gp consists of two homologous halves, each with a transmembrane domain (TMD) followed by a nucleotide-binding domain (NBD), fused in a single polypeptide chain (TMD1–NBD1–TMD2–NBD2) [11]. As documented in structural studies by Rosenberg *et al.* [12, 13], the transporter undergoes significant conformational changes in the presence of nucleotides and substrates. According to the alternative access model [14], the transporter alternates between an inward-facing (IF) conformation, in which the two TMDs form a large internal cavity open to the cytosol and inner membrane leaflet, and an outward-facing (OF) conformation, in which the TMDs rearrange to open toward the extracellular side. It is hypothesized that substrates bind within the IF cavity and are released to the extracellular space upon transition to the OF state, a process energized by ATP binding and hydrolysis at the NBDs [14, 15]. Decades of structural studies have captured multiple inward-facing P-gp structures, all showing a prominent internal cavity capable of accommodating drugs and inhibitors [16, 17, 18, 19, 20, 21]. The first P-gp crystal structure revealed a 6000 Å^3^ solvent-filled cavity open to the cytoplasm and membrane interior, consistent with a “hydrophobic vacuum cleaner” mechanism allowing P-gp to bind a wide variety of compounds [22, 21]. Biochemical analyses confirmed that this cavity can bind at least two substrate molecules simultaneously, indicating multiple ligand subpockets within a common cavity [23, 24]. Indeed, site-directed mutagenesis and photolabeling studies identified at least two primary drug-binding regions, the so-called H site (for Hoechst 33342-like substrates) and R site (for rhodamine 123–like substrates) [25, 23], as well as a third, allosteric site (P site) that can modulate transport [23]. This has led to a model of P-gp as a single flexible drug pocket with partially overlapping binding subpockets. Consistent with this, high-resolution structures of human P-gp have captured ligands bound in different locations within the cavity: for example, a bulky taxane (paclitaxel) was resolved in a pose bridging the two halves of the pocket, contacting helices from both TMD1 and TMD2 [26, 27], whereas smaller inhibitors can occupy more peripheral locations or even bind as pairs in the cavity [26, 28, 17]. In human P-gp, recent structures under turnover conditions reveal a transient peripheral drug-binding site near TM4–TM5 in an inward-open conformation and a distinct central site spanning both halves in more occluded intermediates [17]. These findings reinforce a picture of P-gp ligand cavity as a dynamic, continuum of binding modes rather than a single static site. The capacity of P-gp to bind multiple substrates at once (or one substrate in multiple modes) underlies its polyspecificity, but also raises questions about how such flexible drug binding is coupled to the ATP-driven conformational cycle.

Drug binding and ATP binding/hydrolysis in P-gp are tightly interlinked. Biochemical and biophysical evidence indicate that occupancy of the drug-binding cavity influences the NBDs and vice versa [29, 11, 30, 31]. Early studies showed that transport substrates stimulate ATP hydrolysis activity, suggesting a cooperative dependence of NBD closure on substrate engagement [32, 33, 34, 19, 20]. The two ATP-binding sites of P-gp work in concert and possibly in an alternating manner, both sites are catalytically active but hydrolyze ATP sequentially, with the conformation of one NBD influencing the activity of the other [35, 36]. This coordination implies that substrate binding in one half of the TMD might preferentially stimulate the catalytic site on one NBD, a phenomenon reminiscent of “alternating catalytic sites” behavior observed in kinetic models [37]. There are some evidence for allosteric coupling between the drug-binding TMDs and NBDs, including the hydrogen–deuterium exchange experiments on P-gp, which revealed ligand- and nucleotide-dependent protection changes in the intracellular coupling loops (ICL) that connect TMDs to NBDs, as well as in nucleotide-binding motifs at the NBD interface (H-loop, Walker A), consistent with long-range conformational communication [38, 39]. Complementary molecular dynamics (MD) simulations have shown that a bound drug can promote closer approach and alignment of the NBDs even before ATP is present. For example, in silico studies by Pan and Aller demonstrated that filling the internal cavity with transportable ligands improved the registry of the two NBDs compared to the ligand-free protein, whereas the apo P-gp often closed in a skewed, non-productive orientation [40]. Conversely, nucleotide binding may affect conformational changes in the TMDs that alter the drug-binding pocket [41, 42]. Single-molecule fluorescence microscopy and EPR studies on P-gp in lipid bilayers showed that different drugs bias the ensemble of P-gp toward different conformations [43, 44]. Kinetic and spectroscopic analyses confirmed that verapamil and ATP influence binding of each other: binding of a nonhydrolyzable ATP analog induces distinct P-gp conformations only in the presence of verapamil, and the effects of verapamil on P-gp ATPase switch from activation to inhibition as the nucleotide state or drug stoichiometry changes [45, 46]. Such observations underscore that P-gp functions through a finely tuned allosteric mechanism in which drug occupancy and nucleotide-driven conformational changes are interdependent [47, 45]. Yet, despite these insights, the structural details of how nucleotide binding in the inward-facing state influences the binding mode of a substrate (and vice versa) remain incompletely understood. Key open questions include whether ATP binding can reposition a drug from a peripheral site toward the center of the cavity (as hinted by recent cryo-EM structures), and conversely, whether a bound drug can asymmetrically stabilize one NBD over the other before catalysis as suggested by biochemical asymmetry in the two nucleotide sites of P-gp. Answering these questions is central to understanding how the transport cycle of P-gp is gated by substrate recognition.

Another elusive aspect of the mechanism of P-gp is the role of the long, flexible linker that connects NBD1 and TMD2. This 75-residue linker is present in P-gp and other human ABC transporter isoforms while it is absent in the homodimeric ABC exporters [19]. The region is typically unresolved in crystal and cryo-EM structures due to the high structural mobility, and it was even deleted or truncated in some early structural studies to facilitate crystallization. For a long time the linker was assumed to be simply a non-structured tether with little direct role in transport. However, emerging evidence suggests the linker might modulate the conformational behavior and allostery of P-gp [48, 49, 50, 51]. Biochemical studies have shown that deleting or proteolytically cleaving the linker, while keeping the two halves associated, can alter the ATPase activity of P-gp and drug transport kinetics [52]. The linker contains multiple phosphorylation sites, implicating it in potential regulatory control of P-gp function [53, 54]. Notably, recent enhanced MD simulations of human P-gp indicate that the linker can adopt transient helical structure and engage in interdomain contacts that affect NBD–TMD coupling [51, 50]. Another recent computational study by Barbieri *et al.* simulating various linker conformations in P-gp showed helical arrangement of the linker and associations with other globular P-gp domains [49]. In silico analyses by Ferreira *et al*. suggested that the linker can act as a “molecular damper,” restraining the relative motions of NBD2 and the TMDs by transiently anchoring to the NBD/TMD interface and thus promoting more synchronized domain movements [55]. Consistent with this idea, hydrogen-deuterium exchange experiments detect nucleotide-dependent changes in the regions where the linker and coupling helices converge [38]. Despite being intrinsically disordered, the linker therefore may influence how P-gp transitions from inward-to outward-facing, for example, by biasing the symmetry of NBD engagement or by affecting how the NBDs dimerize which might affect the accessibility of TMD cavity for drug molecules.

The present study seeks to delineate how nucleotide binding and drug binding events are structurally and dynamically coupled in the inward-facing state of human P-glycoprotein. We build upon our previous drug-free simulations of inward-facing P-gp [50] and now introduce transport ligands to directly probe the bidirectional influence between the nucleotide-binding sites (NBSs) and the transmembrane drug-binding cavity. Specifically, we employ extensive all-atom MD simulations of a full-length human P-gp embedded in lipid bilayer to examine P-gp in various nucleotide states (apo, ADP-bound, ATP-bound, and mixed ADP/ATP at the two NBSs) in the presence of representative substrates/drugs. Paclitaxel and verapamil were chosen as model ligands because they are classical P-gp transport probes with contrasting size, rigidity, and effects on ATPase activity (a large, relatively rigid taxane versus a smaller, flexible phenylalkylamine). Each drug was initially docked to distinct subpockets within the inward-facing cavity (the canonical H site, the R site, or a P site) to assess how the binding pose might shift under different nucleotide conditions. By comparing simulations across all combinations of nucleotide states and drug-binding locations, we aim to determine how nucleotide occupancy in the open NBDs influences the positioning and interactions of a bound drug, and reciprocally, how drug engagement in the cavity can affect nucleotide coordination or NBD alignment. Our focus is on the inward-facing conformational landscape preceding the outward-facing transition, as this is the regime in which drugs are captured and prepared for export.

## 2 Results

Full-length models of human P-glycoprotein in IF conformation were constructed and embedded in a POPC/cholesterol membrane environment as the basis for atomistic simulations. Using a standardized protocol [56, 50], we performed all-atom molecular dynamics to characterize the conformational behavior of P-gp in the presence of drug molecule (paclitaxel or verapamil) in five different nucleotide conditions, apo, ADP/ADP, ADP/ATP, ATP/ADP and ATP/ATP; full methodological details are provided in Materials and methods.

### 2.1 Drug–TMD contact residue profile in IF P-gp

In our simulations, paclitaxel and verapamil formed distinct yet overlapping TM contact sets that varied with initial H/R/P docking and nucleotide state. For paclitaxel, the taxane core and aromatic/ester moieties dominate contacts at different helices depending on nucleotide state. For verapamil, dimethoxyphenyl, nitrile, and tertiary amine groups contribute to hydrophobic contacts that shift with nucleotide state and initial docking.

#### 2.1.1 Paclitaxel interaction profiles across nucleotide states and sites

In inward-facing P-gp, paclitaxel engaged a broad set of TM helices with patterns that depended on the initial docking positions in H/R/P sites and nucleotide state (Fig. 1; Supp. Fig. 1). Contact moieties on paclitaxel were the taxane core, aromatic rings (R1–R3), acetate/benzoate esters, and the oxetane ring. Across nucleotide states, paclitaxel maintained site-specific contact profiles dependent on the initial H/R/P docking position. Within each site, nucleotide occupancy altered which paclitaxel moiety accounted for the most persistent contact with a given TM helix.

**Figure 1:**
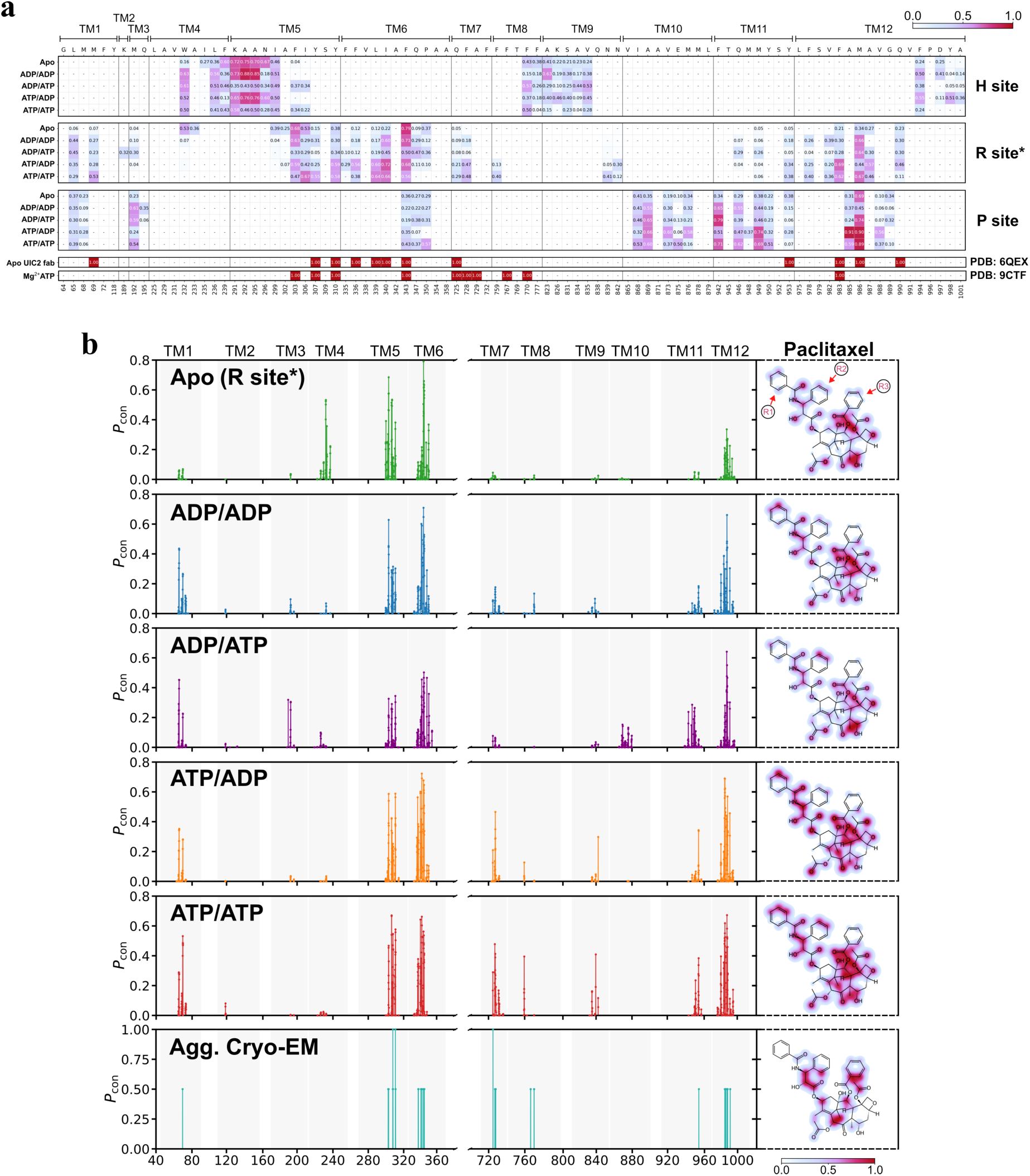
Paclitaxel-TMD interaction profile in IF P-gp. (a) Residue level interaction frequency between the P-gp TMD residues and the bound drug. The frequency from the five nucleotide states are shown for the three P-gp systems with paclitaxel docked at three different sites within TMD, H, R and P site. Previously published cryo-EM structures with paclitaxel bound were used to calculate the same types of interactions [26, 27]. (b) Frequency of atom level interactions between the IF P-gp and the bound drug paclitaxel (TAX) docked initially at the R site, with five nucleotide conditions, apo, ADP/ADP, ADP/ATP, ATP/ADP, and ATP/ATP, each colored in green, blue, purple, orange, and red, respectively. The ligand atoms involved in the interaction were colored with the respective interaction frequencies.

**H-site docked paclitaxel simulations** H-site docked paclitaxel localized near intracellular TM4/5/9/12, with consistent contacts to TM4, TM5, TM8, TM9, and TM12 (Supp. Fig. 1a). Specific ligand moiety-TM helix contacts were observed: taxane core→TM5; acetate/benzoate esters (on the taxane core)→TM4; aromatic ring R1→TM12 in a nucleotide-dependent manner. The subpocket in the H site involved multiple aromatic side chains like W232, F239, F770, F777, and F994 forming hydrophobic and ⇡-stacking interactions with ring moieties of paclitaxel (Supp. Fig. 3). These π-stacking and hydrophobic contacts arose from aromatic rings R1–R3 engaging TM4, TM8, TM9, and TM12. In apo, paclitaxel remained near the H-site subpocket, with high-frequency contacts at TM4 F239 and TM5 (e.g., A292/A295), (Fig. 1). These apo contacts reflected acetate/benzoate esters engaging TM4 near F239 and the taxane core engaging TM5 A292/A295. With nucleotides bound, TM4 W232 contacts became frequent alongside other TM4 residues (Fig. 1). This contact was consistent with π-stacking engagement of aromatic ring moieties of paclitaxel with aromatic residues in TM4. In ADP/ADP and ATP/ADP (NBS2-ADP), paclitaxel remained in the initial H-site subpocket with reinforced TM5 (e.g., A292/A295), TM9 (e.g., A823/K826), and TM12 (e.g., F994/Y998) contacts (Fig. 1a; Supp. Fig. 3c,d). In these NBS2-ADP bound states, the taxane core of paclitaxel accounted for the high TM5 contacts, oxetane ring contributed to contacts with TM9, and aromatic ring R1 contributed to the TM12 contacts at F994 and Y998. In contrast, for the ADP/ATP and ATP/ATP conditions (NBS2-ATP), intracellular TM5 contacts diminished, while center-cavity TM5 residues (F303/I306) became more engaged. This reflected acetate-ester (near the oxetane) engagement of cavity-facing I306 and increased ring-moiety contacts on TM12. In ADP/ATP and ATP/ATP, TM8 F770 contacts increased, while contacts to intracellular TM12 (F994) decreased. In these NBS2-ATP containing states, TM8 engagement was driven by the taxane core of paclitaxel, while contacts with TM12 shifted toward more extracellular residues such as Y998 that were contacted by aromatic ring R1 along with less frequent contact with the intracellular portion of TM12 (F994). Thus, apo and NBS2-ADP favored an intracellular TM4/5 subpocket, whereas NBS2-ATP facilitated subtle displacement toward the central cavity from the H-site.

### R-site docked paclitaxel simulations

In R-site simulations, paclitaxel contacted all inner-cavity helices except TM10, clustering near the cavity center (Fig. 1a). These R site contacts were dominated by the taxane core of paclitaxel engaging the central helices TM6 and TM12 and by additional aromatic ring and acetate ester moieties engaging TM5, TM11, and TM12 in a nucleotide-dependent manner. TM6 F343 at the cavity center was a recurrent top contact in apo and nucleotide-bound P-gp. This prominent TM6 contact reflected direct engagement of the taxane core and aromatic ring R1 moiety of paclitaxel with F343 in the center of the cavity. In apo, contacts concentrated at the center (F343) and the TM4/5 side (e.g., F303/I306), consistent with a slight drift toward the H-site (Fig. 2a). These apo contacts were driven by aromatic ring R1 engaging TM6 residue F343 in the cavity core together with the taxane core and benzoate ester moiety engaging TM5 and adjacent TM4 residues on the same side of the cavity. In ADP/ADP and ADP/ATP (Fig. 1a; Fig. 2b–c), TM1 L65 added hydrophobic contacts while TM6 (I340/F343) and TM12 (M986) remained stably engaged. In NBS1-ADP states, TM6 contact was taxane-core driven; in ADP/ATP condition the taxane also engaged TM12 M986. Notably, ADP/ATP state also promoted contacts to TM2/3 (e.g., K189/M192) near the TM1/P-site region of the cavity. In ATP/ADP and ATP/ATP conditions (NBS1-ATP), paclitaxel shifted upward within the central cavity, engaging upper TM6 (V336/I339), TM7 (F728), TM11 (Y953), and TM12 (F983) via H-bond, hydrophobic, and ⇡-stacking contacts (Fig. 1a–b; Fig. 2d–e). These upper cavity contacts were supported by the taxane core of paclitaxel engaging upper TM6, aromatic ring R3 and benzamide carbonyl engaging TM7, and aromatic ring R1 engaging TM12, together with additional contacts on TM11. With ATP bound on both NBS (ATP/ATP), aromatic moieties further engaged upper TM8, consistent with a higher position toward the extracellular side. NBS1-ATP states (ATP/ADP, ATP/ATP) also added TM7/9 (N839/N842) contacts via the oxetane ring, yielding a denser coordinating network than in NBS1-ADP states, displaying a nucleotide-dependent induced fit phenomenon showcasing the polyspecificity of the dynamic cavity.

**Figure 2:**
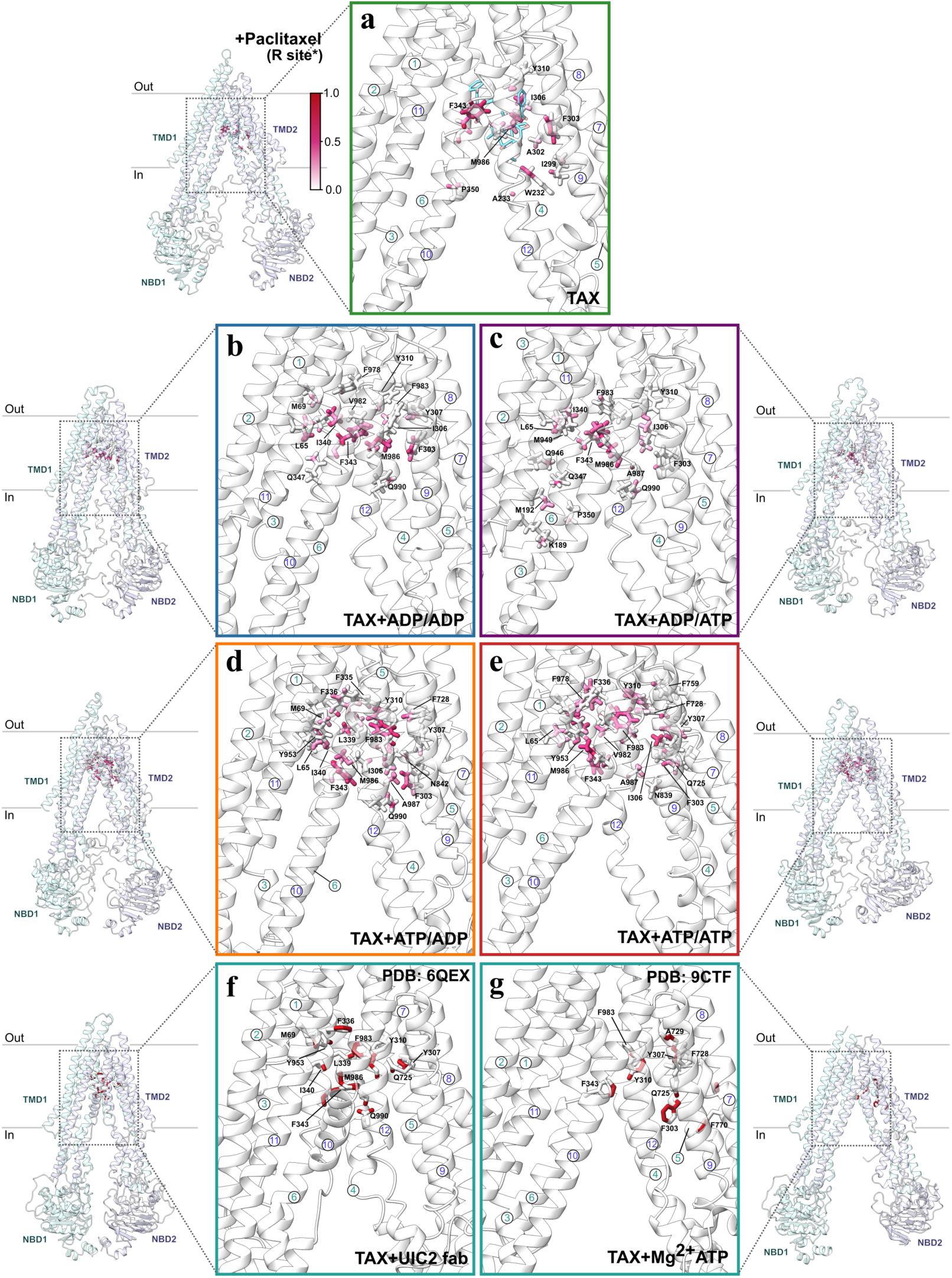
Paclitaxel-TMD interaction profile in IF P-gp. (a-e) Global orientation of IF P-gp shown on the side to display the area within the TMD that is analyzed in each panel. The enlarged picture of the ligand binding site shows residues involved in the interaction with verapamil in nucleotide conditions apo, ADP/ADP, ADP/ATP, ATP/ADP, and ATP/ATP, each in the box colored in green, blue, purple, orange, and red, respectively. Residues with at least one atom with interaction frequency above 0.25 are shown. The initial docking pose of the drug is shown in panel a. (f,g) Interaction profile between the bound paclitaxel and the protein in the cryo-EM P-gp models.

**Figure 3:**
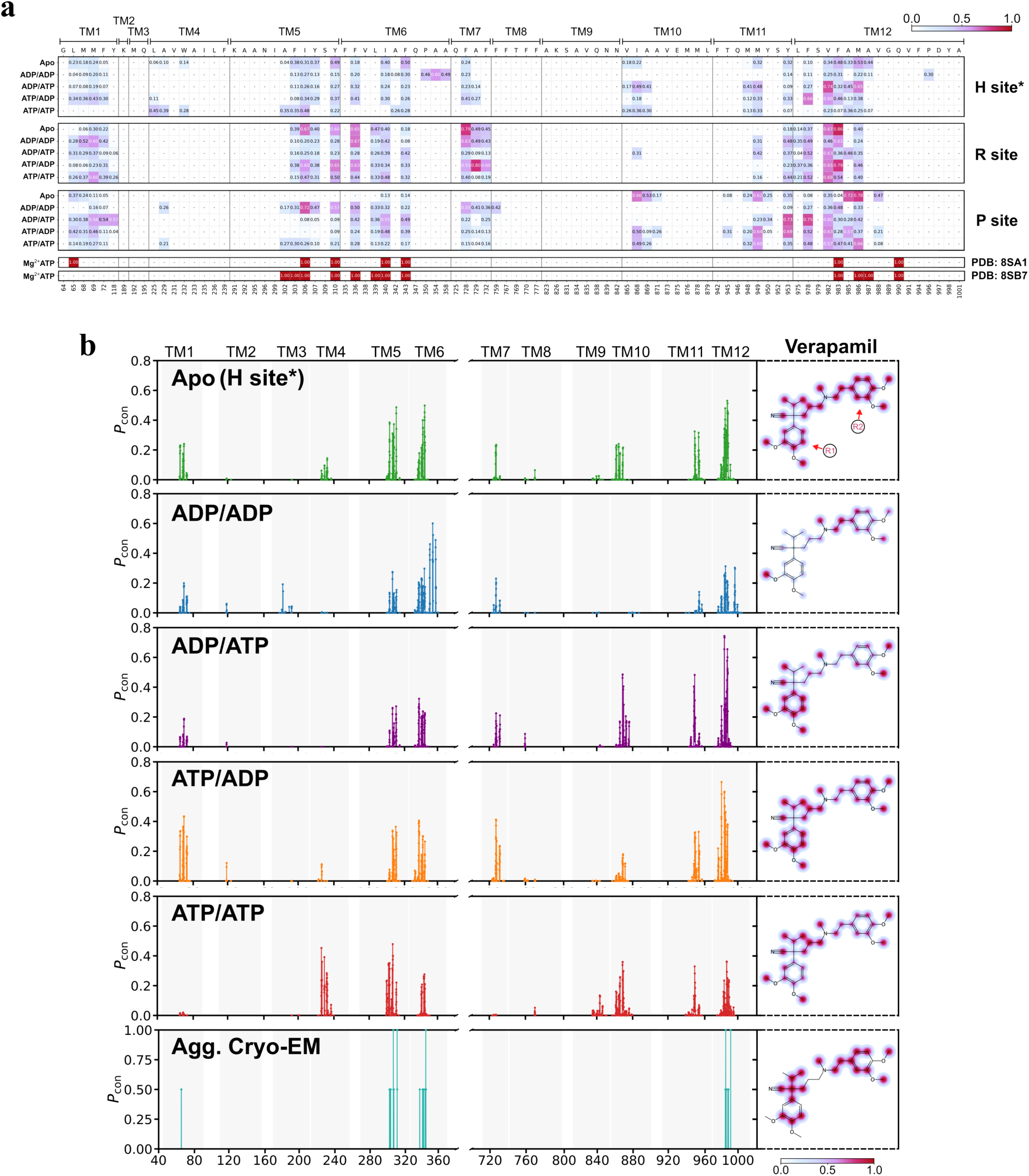
Verapamil-TMD interaction profile in IF P-gp. (a) Residue level interaction frequency between the P-gp TMD residues and the bound drug. The frequency from the five nucleotide states are shown for the three P-gp systems with verapamil docked at three different sites within TMD, H, R and P site. Previously published cryo-EM structures with verapamil bound were used to calculate the same types of interactions [17]. (b) Frequency of atom level interactions between the IF P-gp and the bound drug verapamil (VER) docked initially at the R site, with five nucleotide conditions, apo, ADP/ADP, ADP/ATP, ATP/ADP, and ATP/ATP, each colored in green, blue, purple, orange, and red, respectively. The ligand atoms involved in the interaction were colored with the respective interaction frequencies.

### P-site docked paclitaxel simulations

P site docked paclitaxel displayed consistent interaction near the initial docked region involving TM1, 3, 10 and 11 while contacting the center cavity TM helices 6 and 12 (Fig. 1a, Supp. Fig. 2). These P site contacts arose from defined paclitaxel moieties, where aromatic ring R3 and secondary alcohol near amide moiety engaged TM10, aromatic ring R1 engaged TM11, and the taxane core and ester moieties engaged TM12. In nucleotide-free condition, paclitaxel maintained moderate contacts with all nearby TM helices in the TM1/3 and TM10/11 region closer to the initial docked position. These contacts included engagement of TM1 and TM3 side residues by aromatic ring R2 and benzoate carbonyl of paclitaxel. At the P-site, nucleotide-dependent redistribution was muted relative to the R-site; paclitaxel remained near its initial position with stable TM1/3 (e.g., L65/M192) and TM10/11 (e.g., A869/F942/Q946) contacts. In NBS1-ADP bound states, these TM10 and TM11 contacts were supported by aromatic ring R3 of paclitaxel engaging TM10 residue A869 and aromatic ring R1 engaging TM11 residue F942. With NBS1-ATP (ATP/ADP, ATP/ATP), mid-cavity contacts increased, including TM11 M949 and TM12 A985 (see Fig. 1a). These elevated mid-cavity contacts corresponded to the taxane core of paclitaxel engaging M949 in TM11 and to the benzoate ester moiety engaging A985 in TM12. In NBS1-ATP states (ATP/ADP, ATP/ATP), contacts at the TM10/12 entry (e.g., TM10 V873/E875/M876; TM12 V988) became more frequent. These NBS1-ATP bound conditions recruited specific paclitaxel moieties to the entry site, including aromatic ring R3 engaging V873 (TM10), secondary alcohol near amide moiety engaging E875 (TM10), and ester and taxane regions of the ligand engaging A985 (TM12).

Overall, nucleotide occupancy governed how paclitaxel moieties engaged three regions: an intracellular TM4/5 subpocket that extends to TM9 and TM12 for H-site, a central cavity around TM6 and TM12 for R-site, and a TM10 and TM12 entry region for P-site. TM5 bridged H- and R-site environments, and TM1/3/TM11 bridged P- and R-site contacts, underscoring a continuous binding interface across the cavity.

In summary, across H/R/P docked simulations paclitaxel sampled a continuous binding surface that links the intracellular H-site pocket, the TM6/TM12-centered cavity, and the TM10/12 entry, with nucleotide occupancy determining both position and which ligand moieties dominate contacts. The same helices were stabilized by different parts of the drug depending on state: the rigid taxane core frequently anchored TM5/6/11/12, while aromatic rings (R1–R3) and acetate/benzoate/oxetane groups redistributed engagement toward TM4/8/10/12. At the H site, apo and NBS2-ADP states favored a persistent intracellular TM4/5 subpocket, whereas NBS2-ATP shifted contacts toward the cavity center (greater TM5 F303/I306 and TM8 involvement and a move on TM12 from F994 toward Y998), indicating ATP-dependent displacement from the H pocket. From the R site, paclitaxel remained centrally anchored—recurrently at TM6 F343 and TM12—while ATP in NBS1 drove an upward migration that recruited upper TM6/TM7/TM11/TM12 (and TM8) and added oxetane-mediated contacts to TM7/9, yielding a denser induced-fit network. At the P site, redistribution was more muted: apo and NBS1-ADP stabilized the TM10/11 gate, whereas NBS1-ATP increased mid-cavity TM11/TM12 contacts yet continued to involve the TM10/12 entry, reflecting a mixed entry–cavity pose. TM5 bridged H- and R-site environments and TM1/3/TM11 bridged P- and R-site contacts, underscoring a continuous interface across the cavity. Overall, nucleotide binding acted less to change where paclitaxel binds than to retune how it binds—switching which of the taxane, aromatic, ester, or oxetane groups stabilizes the same helices—with ATP-bound states biasing the ligand toward more central/upper-cavity engagement; this pattern is consistent with paclitaxel-bound cryo-EM models that place the drug in a TM5/6/11/12-lined pocket with additional contacts toward TM7/8 when ATP is present.

#### 2.1.2 Verapamil interaction profiles across nucleotide states and sites

Verapamil, a smaller and more flexible ligand, formed widespread contacts with patterns that varied by nucleotide state yet converged across the H/R/P docked simulations (Fig. 3; Supp. Fig. 2). Dimethoxyphenyl groups (R1/R2) and hydrophobic substituents on the nitrile/tertiary-amine core reweighted contacts across TM helices in varied nucleotide state, rather than a single fixed binding pose. Overall, verapamil engaged a slightly more limited set of key residues per state compared to paclitaxel (on the order of 8–18 residues with >25% interaction frequency; Fig. 3a), consistent with the smaller size of verapamil. Within those residues, individual moieties of verapamil such as dimethoxyphenyl groups could still show stable contacts with a given residue across many frames, indicating that a small number of contact residues could dominate the interaction pattern in each nucleotide state. Unlike paclitaxel, the three docked sites showed a common interacting set: TM1 M69; TM5 I306/Y310; TM6 F336/I340/F343; TM7 F728; TM11 M949/Y953; TM12 F978/V982/F983/M986. At these sites, aromatic rings engaged TM5 I306 and TM12 F983, while dimethoxy termini contacted TM1 M69 and TM12 M986, engaging the same ligand portions. Despite the different initial docked positions, verapamil localized to the central TM6/TM12 cavity and two nearby subpockets toward TM1/11 and TM5/7. The cavity center near TM6 and TM12 was mainly engaged by hydrophobic substituents from the nitrile/tertiary amine core of verapamil, whereas the subpockets toward TM1/11 and toward TM5/7 were predominantly engaged by dimethoxyphenyl moieties that sat against TM1 and TM5.

**H-site docked verapamil simulations**

H-site simulations revealed a nucleotide-dependent shift in verapamil contact patterns, while primarily locating away from the H site and residing in center cavity near R site (Fig. 4). This positional shift of verapamil within the cavity was accompanied by a switch from TM4/5 contacts mediated by the dimethoxyphenyl moieties to contacts in the central cavity where the methyl ends of the dimethoxyphenyl groups engaged TM6 and TM12. Redistribution of the drug position from the initial H site to the center cavity reflected dynamic modes of contact involving different ligand moieties: the nitrile/tertiary-amine core engaged the TM6 interior in some states, whereas aromatic rings engaged TM10/TM11 near the TM10/TM12 entry in others. In apo state (Fig. 4b), contacts shifted away from the initial TM4/5/9 subpocket to the central cavity (TM12 V982–M986) and the TM5/7 side (F303/I306/Y307/Y310), mediated by aromatic moieties, as verapamil moved upward along TM4/5 toward the central cavity. Similar pattern to apo condition was observed in the ATP/ATP state (Fig. 3a; Fig. 4g), where verapamil established TM5/7 contacts (e.g., L225/I306) plus additional TM10/11 and central TM6/TM12 contacts, indicating a distributed binding pattern at the central cavity. In ADP/ADP (Fig. 4c), verapamil shifted laterally toward intracellular TM6/TM12; in apo and ATP/ATP as the R2 dimethoxyphenyl predominantly contacted lower-cavity TM6 (P350/A354/A358) and TM12 (P996). Other moieties engaged minimally, suggesting the obtained binding state represents a loose, lower-cavity intermediate pose in a potential direct entry path from the lipid milieu for hydrophobic ligands like verapamil. Unlike in ADP/ADP condition that showed verapamil contacting lower TM6/TM12, ADP/ATP and ATP/ADP condition led to clustered contacts near central TM12 (e.g., F982/M986), as verapamil distributed furthest into the top portion of central cavity contacting F336 and F978. In these asymmetric nucleotide conditions, the TM12 centered interactions were driven by aromatic ring moieties of verapamil engaging residues such as F982 and M986 together with adjacent dimethoxy groups that stabilized the same region of TM12. In ATP/ADP condition in particular (Fig. 4d), a consistent interaction cluster formed at the top central cavity, indicating nucleotide-dependent positioning. in ATP/ADP, TM1 (L65/M68/M69/F72) and TM7 (F728) contacts increased while TM10 decreased, consistent with a more internal, TM1/TM7 involving interface. These TM1 and TM7 interactions in ATP/ADP were supported by aromatic ring moieties (R1 and R2) of verapamil engaging L65, M69 and F728 while the nitrile/tertiary amine core continued to contact TM12, indicating that distinct parts of the ligand engaged both H and R site sides of the narrowed cavity at the same time.

**Figure 4:**
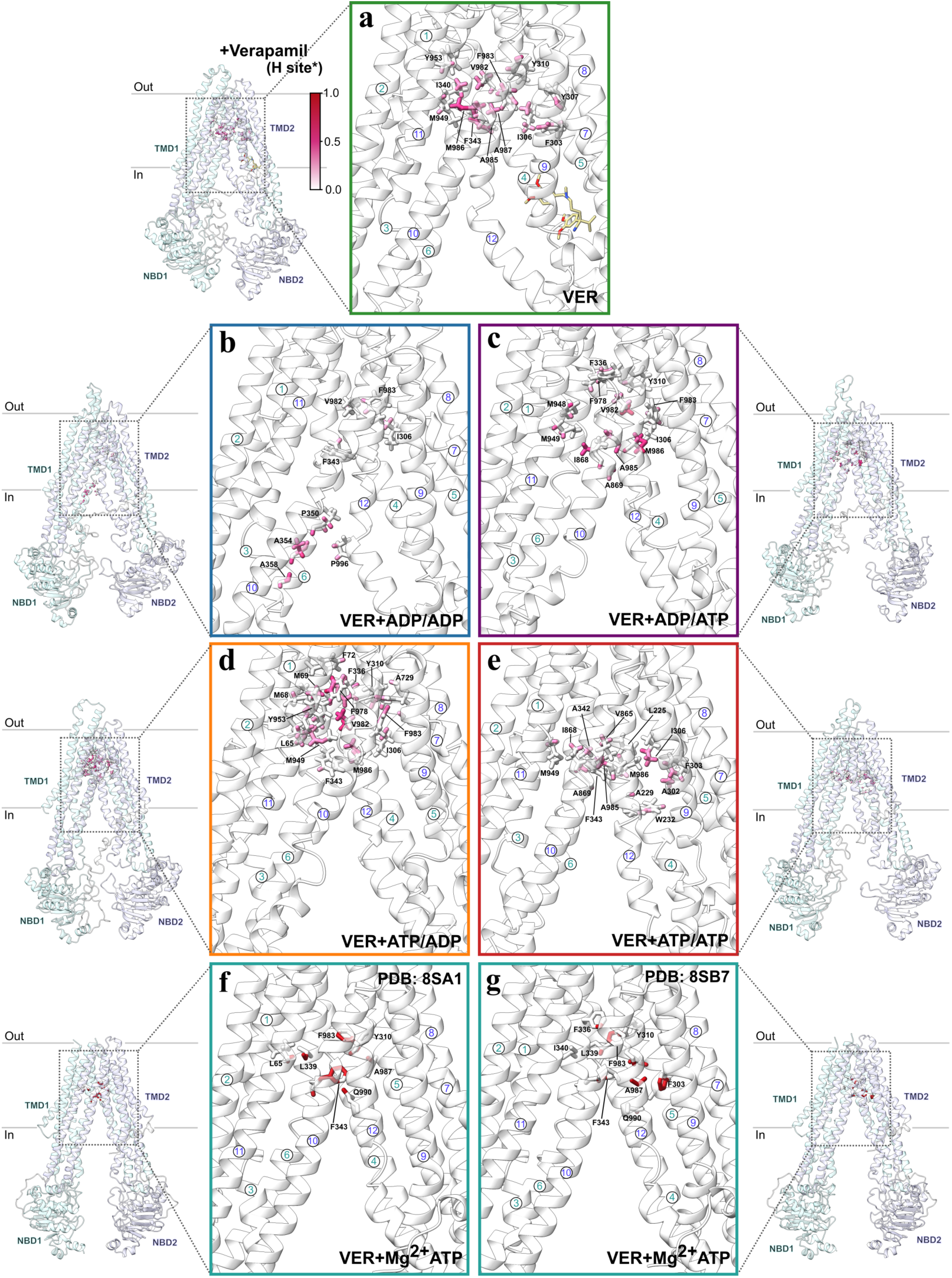
Verapamil-TMD interaction profile in IF P-gp. (a-e) Global orientation of IF P-gp shown on the side to display the area within the TMD that is analyzed in each panel. The enlarged picture of the ligand binding site shows residues involved in the interaction with verapamil in nucleotide conditions apo, ADP/ADP, ADP/ATP, ATP/ADP, and ATP/ATP, each in the box colored in green, blue, purple, orange, and red, respectively. Residues with at least one atom with interaction frequency above 0.25 are shown. The initial docking pose of the drug is shown in panel a. (f,g) Interaction profile between the bound verapamil in the cryo-EM P-gp models.

**R-site docked verapamil simulations**

The R site docked verapamil showed persistent interaction patterns within the central cavity under different nucleotide conditions (Fig. 3a, Supp. Fig. 4a-f). In this R site configuration, the interactions are mediated by almost all parts of verapamil as the drug positions in the central cavity near TM6/12. The apo state simulation results in higher frequency interactions at the central/TM5 side of the cavity with residues I306 (TM5) 6̃7, F728 (TM7) 7̃9 and F983 (TM12) 8̃6%, where I306 on TM5 and F983 on TM12 are engaged by the aromatic rings and the nitrile/tertiary amine core of verapamil. ATP/ADP condition shows a simliar pattern of interactions with I306 ̃60, A729 ̃80 and F983 ̃79%. In ATP/ADP, I306 on TM5 is contacted by both the aromatic ring R1 moiety and dimethoxy moiety of verapamil, indicating that multiple chemical groups of the ligand reinforce the same TM5 interface while the TM7 and TM12 contacts remain strong. While apo and ATP/ADP conditions showed cluster of residues interaction with verapamil in central cavity closer to TM5/7, ADP/ADP, ADP/ATP and ATP/ATP conditions showed slightly skewed interaction profile in central cavity toward TM1. In ADP/ATP condition, verapamil interacted with L65 ̃34, M68 ̃29, and M69 ̃37%. Notably, these TM1 contacts are observed more frequently in ADP/ADP (M68 5̃2, M69 6̃0%) and ATP/ATP (M68 ~ 37, M69 6̃0%). In addition, upper cavity contacts on TM1 (further up in the cavity toward the extracellular side) are established involving F72 (4̃2 in ADP/ADP and 3̃9% in ATP/ATP) and Y118 (2̃6% in ATP/ATP), facilitated by dimethoxyphenyl groups of verapamil rotating toward the extracellular direction. Overall, verapamil initially docked in R site near TM5, 6, 7 and 12 maintained the position within the center cavity in apo and all nucleotide states (Supp. Fig. 4a-f). Although this position is preserved, nucleotide asymmetry shifts which moiety of verapamil provides the dominant anchor, with aromatic ring groups engaging one side of the cavity in some states and other substituent groups engaging the opposite side in others. The contact residues shows that verapamil does not move away from the initial position in apo and ATP/ADP conditions, whereas ADP/ATP condition favors verapamil to interact on the other side of the cavity compared to the initial position, involving TM1. This TM1 engagement in ADP/ATP arises from reorientation of a dimethoxyphenyl group of verapamil toward TM1 while the rest of the ligand remains in the central cavity. In ADP/ADP condition, verapamil moves further up in the center cavity as it forms a stable interaction pattern with residues further up toward extracellular side. Similarly, the ATP/ATP condition favors verapamil to move up in the cavity, but the contact residues are distributed away from TM7, closer to TM1 side of the top cavity. In ATP/ATP this redistribution toward the TM1 side does not rely on one single ligand moiety, and instead multiple moieties of verapamil contribute moderate contacts across TM1 side residues, displaying a possible preferential binding region when favorable nucleotides are bound in the NBDs.

**P-site docked verapamil simulations**

P-site verapamil also concentrated near central TM6/TM12, with nucleotide-dependent changes that differed from the R-site (Supp. Fig. 4g–l). At the P-site, nucleotide states favored different verapamil moieties, with dimethoxyphenyl moieties stabilizing either the TM10/TM12 gate or the TM4/TM5 side. In apo, frequent contacts (e.g., TM1 L65; TM10 I868; TM11 M939; TM12 A985/M986) matched the initial TM1/10/12 subpocket. These TM10 and TM12 contacts in the apo state involved the same dimethoxyphenyl moieties of verapamil that stabilized the TM10 and TM12 gate region in nucleotide bound states. With nucleotides bound, contacts extended to TM5/6/7, redistributing further into the cavity away from the TM10/12 entry vicinity. In ADP/ADP, TM10 contacts diminished relative to apo, while TM5/6/7 contacts (e.g., TM5 I306/Y310; TM6 F336; TM7 F728) increased. In this ADP/ADP state the aromatic ring R2 moiety of verapamil shifted toward TM5 and TM6 to stabilize I306 and F336 contact while TM10 engagement was reduced. In ATP/ATP, contacts distributed between the TM5 side (e.g., F303) and the central cavity (TM11 M949; TM12 F983/M986). These TM5 and central cavity interactions were supported by several different moieties of verapamil, including the aromatic ring R2, a nitrile substituent, and the tertiary amine core, consistent with a broad engagement pattern. In ADP/ATP and ATP/ADP, verapamil engaged the cavity center (TM6/TM12) and the TM1 side of the cavity (e.g., M69/F72/Y118 in ADP/ATP; L65/M69 in ATP/ADP). In these asymmetric nucleotide states, the cavity center contacts on TM6 and TM12 were maintained by the dimethoxyphenyl moiety on aromatic ring R2 side of verapamil, while the TM1 side contacts such as M69 and F72 arose from the aromatic ring R1 that rotated toward TM1. Verapamil remained near the TM10/12 entry in apo and, to a lesser extent, in ATP/ADP. In ATP/ADP, both the aromatic ring and the methoxy portions of verapamil continued to engage TM10 and TM12 in that same entry region, consistent with minimal displacement of the ligand body. In ADP/ATP, verapamil moved away from TM10 along TM1/TM12 to a higher cavity position; in ATP/ATP, it showed a distributed central-cavity pattern involving TM1, 4, 5, 6, 7, 10, 11 and 12.

In summary, many of the H/R/P site docked verapamil contacts shared similar TM residues near the cavity center, implying that the binding modes of verapamil might have interconverted as the conformation of the protein shifted in the presence of different nucleotides in NBDs. The same central residues on TM6 and TM12 could be engaged either by aromatic ring moieties or by methoxy moieties of verapamil depending on nucleotide state, suggesting that changes in nucleotide state could reshuffle which chemical group of the ligand was used to stabilize the same region of the cavity. Interestingly, verapamil did not favor the initial H site docked position in all nucleotide conditions, as most nucleotide states favored verapamil interacting further up in the cavity away from both the TM4/5 and TM10/12 entry gate. This relocation was accompanied by a shift from contacts at TM4 and TM5 that involve both aromatic ring and methoxy moieties to deeper contacts where the aromatic ring moiety engaged TM1, TM7, and TM12 in the central cavity. Exceptions: H-site ADP/ADP and P-site apo both favored residence near the TM10/12 entry. In both of these cases, the TM10 and TM12 entry region was stabilized by aromatic ring and dimethoxy groups of verapamil that remained positioned near the gate rather than shifting deeper into the cavity. Notably, ATP/ATP across H/R/P yielded a broadly distributed binding pattern, consistent with a central cavity that accommodated multiple verapamil poses across inner helices. In ATP/ATP, simultaneous moderate engagement of multiple helices involved dimethoxyphenyl, nitrile, and tertiary-amine groups rather than a single dominant interaction.

### 2.2 CAVER tunnel analysis in IF P-gp with paclitaxel and verapamil

CAVER identified distinct tunnels linking the cytosol and lower lipid leaflet to the ligand-binding cavity in IF P-gp (Fig. 5). Two IF conformers with distinct linker states were analyzed, IF-mdl (unstructured linker) and IF-af (α-helical residues 679–688), and the position of the linker between the two halves modulated which tunnels connected the cytosol/lower leaflet to the cavity. Tunnel definitions: 1, intracellular opening; 2a, TM4/6 from intracellular solvent; 2b, TM4/6 from lower leaflet; 2c, TM10/12 (intracellular solvent or lower leaflet); 3, inner sub-pocket near TM4/7/8.

**Figure 5:**
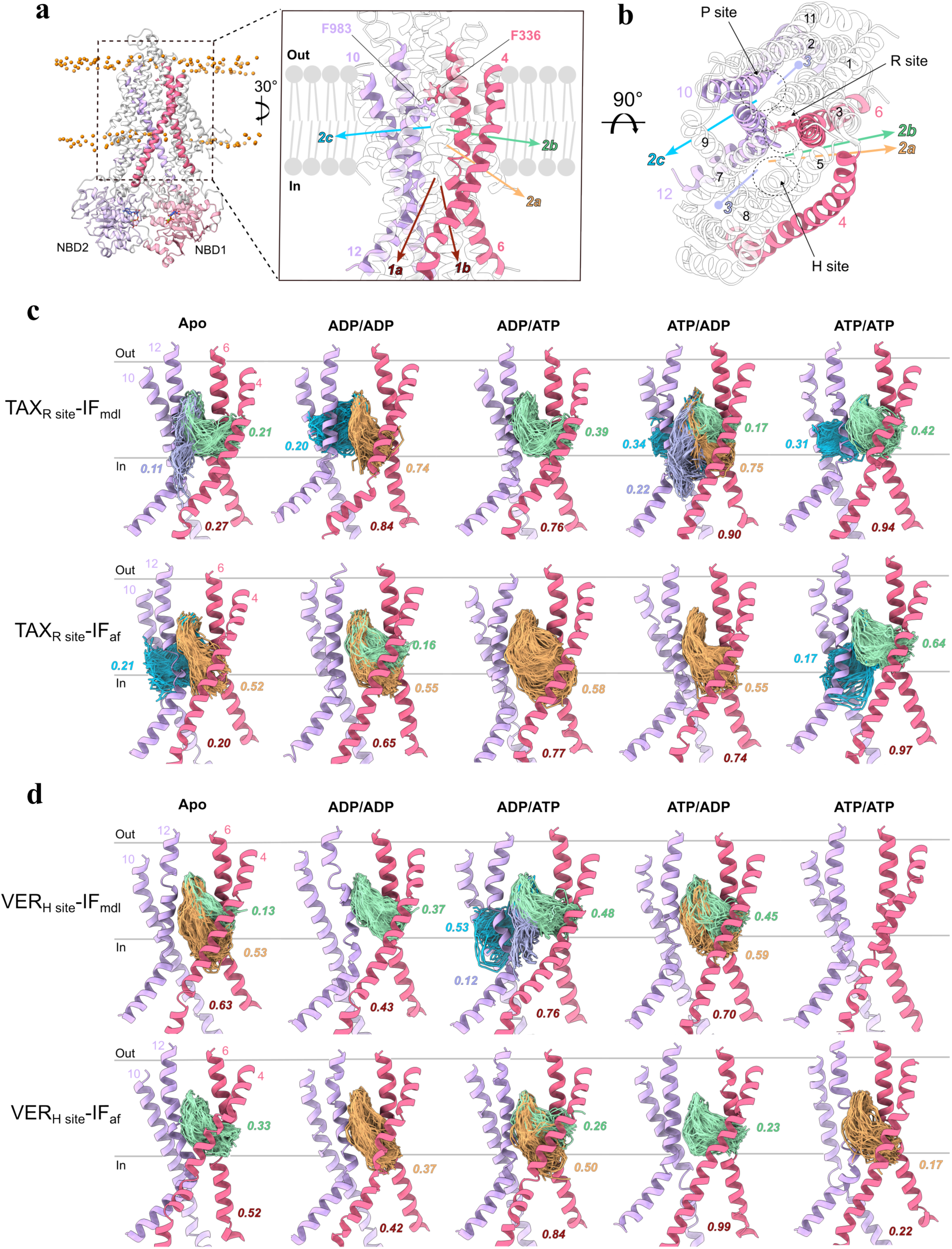
CAVER-identified ligand-accessible tunnels in IF P-gp with paclitaxel (TAX) or verapamil (VER). Probes were initiated at F336 and F983. Tunnel types include 1 as the intracellular opening, 2a and 2b as TM4/6 routes from intracellular solvent and lower leaflet, 2c as TM10/12, and 3 as an inner sub-pocket near TM4/7/8. Occupancies were normalized per replica; the last 70% of frames were analyzed; only tunnels with occupancy greater than 0.10 are shown. Panels a and b show views of identified tunnels, panel c shows TAX, and panel d shows VER.

**Figure 6:**
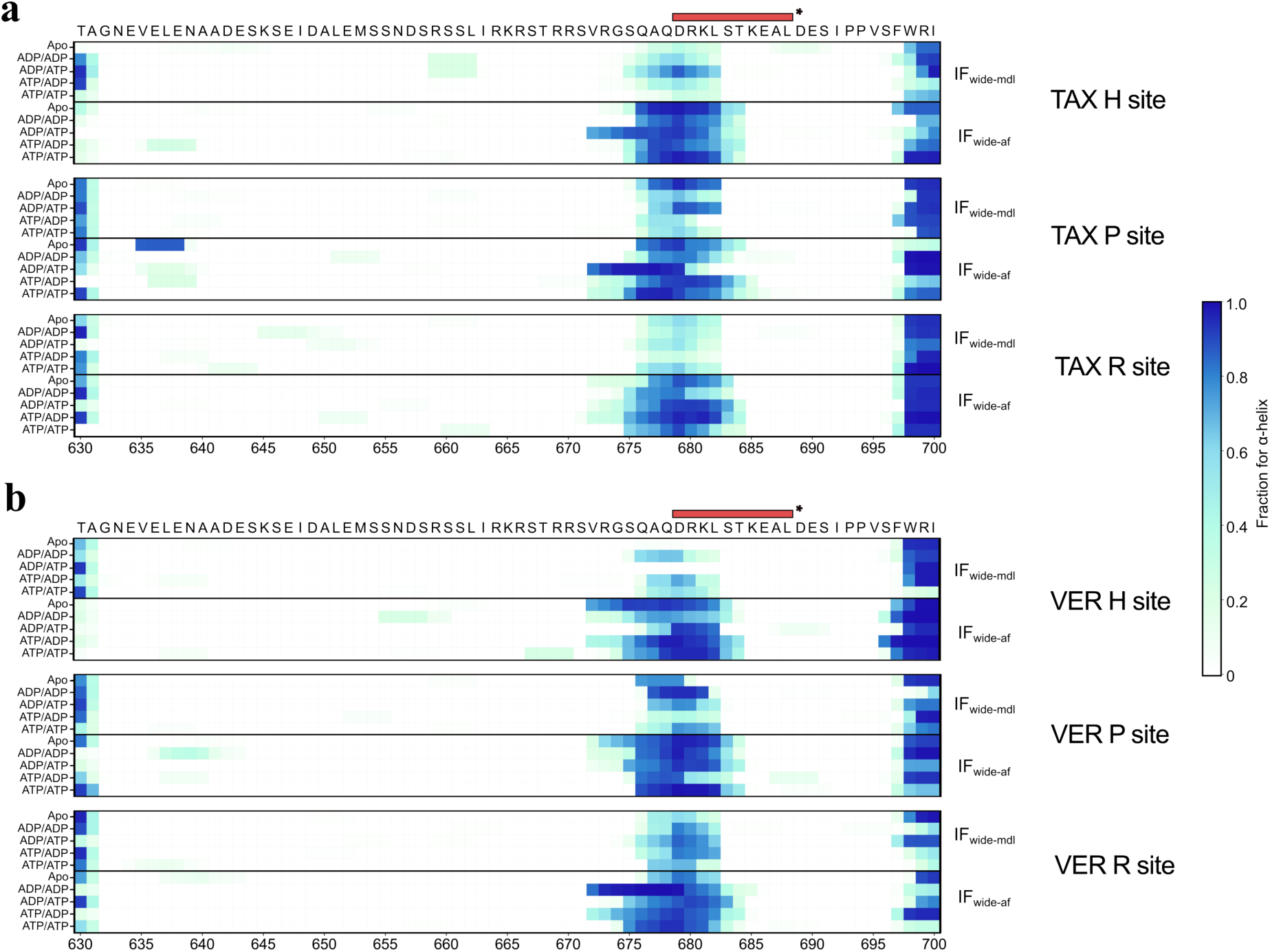
Flexible linker helicity in IF P-gp with paclitaxel (TAX) or verapamil (VER) across five nucleotide states: apo, ADP/ADP (NBS1-ADP/NBS2-ADP), ADP/ATP (NBS1-ADP/NBS2-ATP), ATP/ADP (NBS1-ATP/NBS2-ADP), and ATP/ATP (NBS1-ATP/NBS2-ATP). Helicity for residues 630–700 is normalized over replica trajectories and shown as heatmaps for IF-mdl and IF-af. The red bar marks the AF2 helix (679–688) used in the IF-af template. (a) TAX. (b) VER.

#### Ligand pathways observed in paclitaxel-bound IF P-gp

For R-site paclitaxel, channel1 was the most frequent tunnel across ADP/ADP, ADP/ATP (NBS1-ADP/NBS2-ATP), ATP/ADP (NBS1-ATP/NBS2-ADP), and ATP/ATP (NBS1-ATP/NBS2-ATP). In IF-mdl, channel1 occurred in 84% (ADP/ADP), 76% (ADP/ATP), 90% (ATP/ADP), and 94% (ATP/ATP) (Fig. 5c). Channel2a was next most frequent in IF-mdl, with 73% (ADP/ADP) and 75% (ATP/ADP). By contrast, the apo IF-mdl state showed much lower tunnel frequencies (channel 1 in only 27%; channel 2a 21%; channel 3 11%). In IF-af, channel1 was likewise frequent: 64% (ADP/ADP), 76% (ADP/ATP), 74% (ATP/ADP), 96% (ATP/ATP). Channel2a was generally second most common (54–58%) across apo and all nucleotide states except ATP/ATP. In IF-af apo, channel2a was 52%, channel1 was 19%, and channel2c was 20%. In IF-mdl, ATP/ADP yielded concurrent openings in multiple tunnels (1: 90%, 2a: 75%, 2b: 17%, 2c: 34%, 3: 22%). For channel2c (TM10/12), IF-mdl showed 2c in all states except ADP/ATP, whereas IF-af showed 20% only in apo and ATP/ATP. Channel occupancies depended on the nucleotide at NBS1 versus NBS2. With NBS1-ATP (ATP/ADP, ATP/ATP), channel1 was 90–94% (IF-mdl) and 74–97% (IF-af), versus 76–84% (IF-mdl) and 65–77% (IF-af) with NBS1-ADP. In IF-mdl, channel2a appeared only with NBS1-ADP (ADP/ADP, ADP/ATP).

Across H and P site paclitaxel, channel1 was usually most frequent across ADP/ADP, ADP/ATP, ATP/ADP, and ATP/ATP, with patterns modulated by the docked site and linker conformation. In IF-mdl (TAX-H), channel1 occurred in 77% (apo, ADP/ADP), 84% (ADP/ATP), 66% (ATP/ADP), and 88% (ATP/ATP). In IF-mdl (TAX-H), channel2a ranged from 30% (ATP/ADP) to 62% (ATP/ATP) and was absent in ADP/ATP; channel2b reached 45% (ATP/ATP), and channel2c reached 45% (ADP/ADP), providing nucleotide-dependent alternatives to channel1. The IF-mdl with paclitaxel docked at P site (TAX-P site) showed a contrasting profile: channel 1 was modest in apo (26%) and only 51% in ADP/ATP, 36% in ATP/ADP, yet higher in ADP/ADP (83%) and ATP/ATP (79%). When channel1 was low in TAX-P (e.g., ADP/ATP or ATP/ADP), channels2a/2b/2c became more prevalent (ADP/ATP: 27/27/16%; ATP/ADP: 2c28%). Overall in IF-mdl, TAX-H maintained channel1 at 66–88% across states, whereas TAX-P showed low channel1 in apo (26%), ADP/ATP (51%), and ATP/ADP (36%), with channels2a/2b/2c each contributing 16–28% in those cases.

IF-af showed a different distribution than IF-mdl. In the H site docked simulations, channel 1 appeared in 79% of apo, 30% of ADP/ADP, 80% of ADP/ATP, 52% of ATP/ADP and 89% of ATP/ATP frames. In ADP/ADP condition for the H site docked IF-af, channel 1 was 30% while channel 2b was 21%. In the P site docked IF-af, channel 1 was higher in apo (93%) but again low in ADP/ADP (45%), and moderate in ADP/ATP (54%), ATP/ADP (49%) and ATP/ATP (71%). In IF-af, channel2b was frequent (H: 43% apo, 53% ATP/ATP; P: 29% apo, 29% ATP/ADP), and at the H site channel2a reached 52% (apo) and 57% (ADP/ATP) but was 37% in ATP/ATP. With ATP/ATP, channel1 was high in both conformers (IF-mdl: R 94%, H 88%, P 79%; IF-af: R 96%, H 89%, P 71%), whereas in ADP/ADP IF-af dropped to 30% (H) and 45% (P).

#### Ligand pathways observed in verapamil-bound IF P-gp

Verapamil sampled multiple tunnels with frequencies dependent on the docked site, NBS state, and IF-mdl versus IF-af. At the H site, channel1 dominated in IF-af across all NBS states and in IF-mdl for all but ATP/ATP, with state-dependent magnitudes. In IF-mdl, channel 1 occurred in 63% of apo simulations, decreased to 42% in ADP/ADP, then increased to 76% in ADP/ATP and 78% in ATP/ADP. In IF-mdl ATP/ATP, no tunnels (including channel1) were detected. In IF-af, channel 1 occurred at 51% in apo and 41% in ADP/ADP, increased to 83% in ADP/ATP and 98% in ATP/ADP, and then decreased to 22% in ATP/ATP. Channel2a showed opposite trends in IF-mdl versus IF-af across states. In IF-mdl, channel 2a was present at the H site in apo (52%), ADP/ADP (37%), ADP/ATP (47%), and ATP/ADP (58%), and was not detected in ATP/ATP. By contrast, in IF-af at the H site, channel 2a reached 49% in ADP/ATP, was 32% in apo and 37% in ADP/ADP, and then decreased to 22% in ATP/ADP and 16% in ATP/ATP. At the H site, channel2b occurred in IF-mdl at 13% (apo) and 45% (ATP/ADP) but not in ADP/ADP, ADP/ATP, or ATP/ATP; in IF-af it appeared only in ADP/ATP (26%). Channel2c was limited to IF-mdl (53% in ADP/ATP) and absent in IF-af. Finally, a less frequent channel 3 emerged in IF-mdl under ADP/ATP (11%) but was absent in IF-af simulations. Thus, at the H site, channel1 was frequent in nearly all states, channel2a was common in apo, ADP/ADP, ADP/ATP, and ATP/ADP, and channels2c (53%) and3 (11%) appeared only in IF-mdl ADP/ATP. In ATP/ATP at the H site, IF-af retained channel1 (22%) whereas IF-mdl showed no tunnels; IF-af lacked channels2c/3, showing only 2a and, in ADP/ATP, 2b. Uniquely, IF-mdl ATP/ATP at the H site showed no tunnels, whereas other states had at least one. At the H site in ATP/ATP, IF-af retained channel1 (22%) and 2a (16%), confirming that the ATP/ATP occlusion was specific to IF-mdl.

For P- and R-site verapamil, channel1 was typically observed the most, with magnitudes and secondary tunnels varying by site, NBS state, and IF-mdl versus IF-af. In the apo condition, channel 1 frequency at the R site was 87% in IF-mdl and 91% in IF-af, compared with 51% in IF-mdl and 64% in IF-af at the P site. In IF-mdl ADP/ADP, P-site showed channel2c at 65% (absent at R-site), whereas R-site showed 2b at 54% and 2a at 12%. In IF-af ADP/ADP, P-site showed channel2a at 40% versus 16% at R-site; channel2b occurred at both (25% vs 43%). In ADP/ATP, IF-mdl maintained channel1 (P 82%, R 88%) with similar channel2b (0%). In IF-af ADP/ATP, channel1 stayed high at P-site (93%) but dropped to 58% at R-site, where channel2b matched it (58%). In the ATP/ADP state, channel 1 reached 96% at the P site and 99% at the R site in IF-mdl, and was also frequent in IF-af at 72% for the P site and 84% for the R site. In ATP/ATP conditions, there was a notable difference between the two P-gp conformers: IF-mdl retained high channel 1 (74% at P, 89% at R) with multiple auxiliary tunnels (channel 2c 48% at P, channel 3 29% at R). By contrast, in ATP/ATP IF-af, channel 1 at the P site decreased to 27% whereas at the R site it remained at 81%. In IF-mdl, R-site verapamil consistently sampled channel3 (apo 21%, ATP/ADP 25%, ATP/ATP 29%), which was less frequent or absent at P-site. Overall, R-site verapamil preserved channel1 across states in both conformers, whereas P-site verapamil reduced channel1 specifically in ATP/ATP, especially in IF-af. Under ATP-containing states, IF-mdl sampled more auxiliary tunnels (2b/2c/3), whereas IF-af showed a P-site-specific reduction of channel1 in ATP/ATP (27%) while R-site remained 81%. These trends indicated that, under identical NBS conditions, the initial binding site and linker conformation shaped cavity-accessible tunnel formation (Fig. 5d; Supp.Fig. 5c).

### 2.3 Linker region α-helicity in IF P-gp

Across all IF P-gp simulations, the NBD1–TMD2 linker (residues 630–699) was highly flexible. A transient α-helix repeatedly formed in a central segment (675–685), with occupancy varying by system. IF-af (AlphaFold2 [57]; initial 679–688 helix) generally retained a helical segment, whereas IF-mdl (Modeller [58]; initially unstructured) showed lower occupancy but frequent helix formation under specific nucleotide states. In apo runs (no nucleotide at either NBS), H-site docking of either paclitaxel or verapamil gave IF-mdl with moderate/low helix at Q676–L682 (<0.5), whereas IF-af showed a consistent helix in the same region. In contrast, P-site docking of either drug produced a helix in both IF-mdl and IF-af, suggesting P-site occupancy biased the flexible linker toward an α-helical arrangement.

Nucleotide state and drug identity influenced linker helicity in both IF models. With TAX at the H or P site, ADP/ATP (NBS1-ADP/NBS2-ATP) increased helix occupancy versus apo in IF-mdl, and in IF-af extended the helix toward the N-terminus (V672–G674). In TAX–H IF-mdl, ADP/ATP yielded Q676–L682 0.5 (0.53–0.86; peak D679 0.86). In TAX–H IF-af, ADP/ATP extended helicity to V672–G674 (0.72–0.83), whereas 676–681 were already high across states. At the P site, ADP/ATP similarly gave IF-mdl with 679–682 0.8 and IF-af with a stable N-terminal-shifted helix at 672–679 (0.9). In ADP/ADP, ATP/ADP, and ATP/ATP, TAX-H and TAX-P showed low 676–682 helicity in IF-mdl, whereas IF-af retained high helicity at 675–685. For TAX–R IF-af, ATP/ATP (NBS1-ATP/NBS2-ATP) lowered 675–684 helicity (only Q678/D679 0.8), indicating unwinding of the initial AF helix (679–688) when TAX occupied the top cavity (Fig. 2e). In VER-bound IF-mdl, 676–682 helicity lacked a consistent pattern across nucleotide states, unlike TAX-bound IF-mdl. Notably, VER–H IF-mdl showed no helicity at 670–690 in apo and in ADP/ATP (NBS1-ADP/NBS2-ATP), contrasting with the helicity observed in other TAX/VER IF-mdl and IF-af cases. In VER–H (apo and ADP/ATP), contacts centered on TM1, TM5, TM6, TM10, TM11, and TM12 (Fig. 4a,c), consistent with a non-helical linker and a tighter NBD approach. In VER–H IF-af, ADP/ADP reduced 676–682 helicity (<0.7) relative to other IF-af states. In VER–H ADP/ADP, the ligand localized near the TM10/12 entry (Fig. 4c), suggesting entry-gate engagement could reduce linker helicity. For VER–P, IF-af showed consistent 676–683 helicity except in ATP/ADP (NBS1-ATP/NBS2-ADP), which maintained helicity only at 676–679 (0.73–0.82). VER–P IF-mdl showed a similar trend: ATP/ADP reduced 677–683 helicity (<0.3) relative to other states (0.3–0.95). Under ADP/ATP (NBS1-ADP/NBS2-ATP), verapamil adopted a stable pose contacting TM1, TM6, TM10, TM11, and TM12 near the TM1 and TM10 side of the cavity (Fig.S3.4i). This suppression correlated with stable drug–TMD contacts. Conversely, VER–R IF-af in ADP/ADP displayed an N-terminal-shifted helix at 672–683 while the ligand contacted a tight cluster at the cavity top.

Overall, α-helicity localized to a single cluster within the linker. When present, the helix centered on Q676–L682 (often ±1–2 residues), indicating a context-dependent helical tendency in this segment. Consistently, residues 630–670 showed 0.1 helicity except in apo TAX–P IF-af, where V635–E639 reached 0.78. Some residues within 676–682 had been proposed as phosphorylation sites; here we reported helicity only. S667 and S671 were 0.1, whereas S661 formed a single-turn helix in TAX–H IF-mdl under ADP/ADP and ADP/ATP (0.35) and in TAX–R IF-af (0.14). S675 and S683, within/adjacent to 676–682, were occasionally incorporated into the transient helix. In summary, across initial models, all IF simulations showed a variable linker helix centered at 676–682. The extent of this helicity was modulated by nucleotide state and by ligand identity and initial binding pose. The linker fluctuated between coil and α-helix, consistent with intrinsic disorder reported for this region.

### 2.4 Interaction frequencies between nucleotides and NBS in drug-bound IF P-gp Paclitaxel bound IF P-gp: NBS residue–nucleotide interaction profiles

With paclitaxel, NBS2-bound nucleotides showed lower overall contact frequencies than NBS1-bound nucleotides, especially at the adenosine–ribose region (Fig. 7b,c). This indicated looser coordination as measured by contact frequency at NBS2 than NBS1, which might have favored replacement of NBS2-ADP by ATP during the transport cycle. NBS1-bound nucleotides displayed more consistent contacts overall, yet phosphate groups maintained persistent contacts in both NBS1 and NBS2 across most ATP/ADP states. However, a few conditions showed weaker phosphate coordination than the corresponding states. For instance, TAX P-site IF-mdl showed inconsistent coordination of ADP α and β-phosphate groups at NBS2 in ADP/ADP condition and NBS1 in ATP/ATP condition. Similarly, TAX-H site IF-af showed inconsistent α/β-phosphate coordination for NBS1-ATP in ATP/ATP, and TAX-P site IF-af showed inconsistent α/β-phosphate coordination for NBS2-ADP in ADP/ADP. TAX-P site IF-af also showed less frequent β- and γ-phosphate contacts for NBS2-ATP in ADP/ATP. Atom-level contact frequencies indicated that non-canonical paclitaxel contacts near H/P sites reduced phosphate coordination, suggesting drug–cavity interactions could destabilize NBS-bound nucleotides. Such destabilization likely promoted exchange between NBS-bound and solvent nucleotides, particularly replacement of NBS-ADP by ATP during cycling.

**Figure 7:**
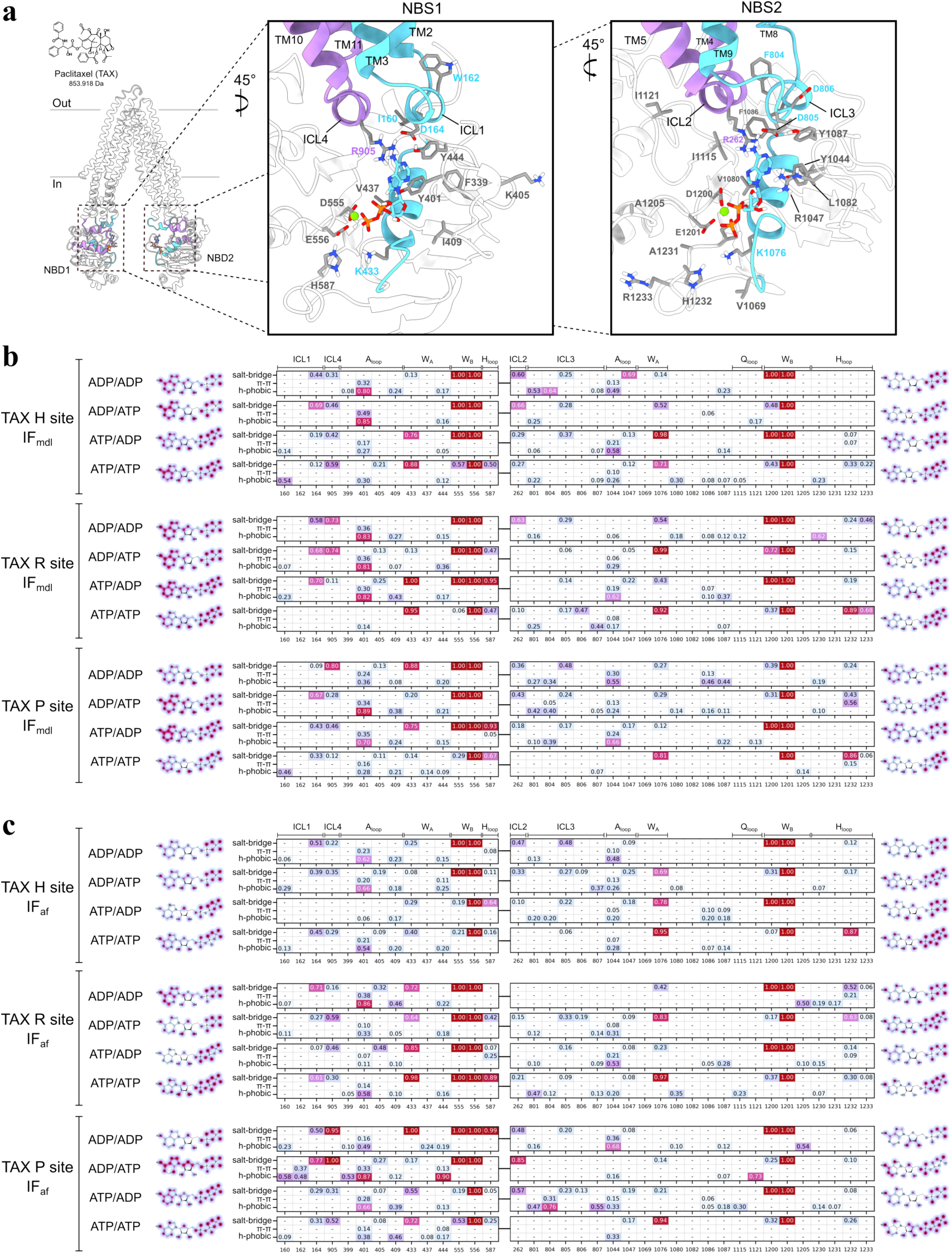
Paclitaxel-bound IF P-gp alters NBS–nucleotide coordination. Interaction profiles between bound nucleotides and NBS residues were calculated. (a) NBS1 and NBS2 residues are labeled; ICL1/3 and Walker A are cyan; ICL2/4 are purple. (b) Heatmaps for TAX IF-mdl show salt-bridge, ⇡-stacking, and hydrophobic interaction frequencies (rows = interaction types; columns = residues), normalized over replicas. Blocks correspond to TAX-H, TAX-R, and TAX-P IF-mdl, each with four nucleotide conditions (ADP/ADP, ADP/ATP, ATP/ADP, ATP/ATP). Adjacent 2D drawings contour nucleotide atoms by interaction frequency. (c) Same format for TAX IF-af.

Numerous conserved NBS1 and NBS2 residues in the inward-facing P-gp remained closely associated with the bound nucleotides (Fig. 7a). Interaction frequencies (Fig. 7b,c) showed Walker A lysines (K433, K1076) and Walker B D/E residues (D555/E556; D1200/E1201) forming phosphate salt bridges across conditions, often at high occupancy. Walker B glutamates showed near-constant nucleotide contact across simulations, whereas Walker B aspartates varied widely (0.06–1.0). In TAX-R systems, Walker A–phosphate contacts were stronger at ATP-bearing sites (NBS1 in ATP/ADP; NBS1 and NBS2 in ATP/ATP) than at ADP-bearing sites (NBS1 in ADP/ADP; NBS2 in ADP/ATP) in both IF-mdl and IF-af. By contrast, TAX-H and TAX-P showed no clear difference in Walker A coordination between ATP-bearing and ADP-bearing NBS. Thus, central (R-site) positioning appeared to favor tighter Walker A coordination for ATP than for ADP. H-loop histidines (H587, NBS1; H1232, NBS2) also contacted nucleotides, with variability across nucleotide state and docked site. These histidines coordinated nucleotide phosphates but without a consistent nucleotide- or drug-dependent pattern. An asymmetry was evident: NBS1 contacts involved H587 alone, whereas NBS2 contacts involved H1232 together with adjacent H-loop residues I1230, A1231, and R1233.

The intracellular coupling loop (ICL) frequently engaged the nucleotide nitrogenous base in several conditions. In the NBS1 region, D164 (ICL1) and R905 (ICL4) exhibited salt bridge contacts in high frequency with the bound nucleotide. In TAX-R site IF-mdl, nucleotide conditions with ADP bound in NBS1 resulted in consistent contacts with nucleotide adenine for D164 (0.58 in ADP/ADP; 0.68 in ADP/ATP) and R905 (0.73 in ADP/ADP; 0.74 in ADP/ATP), whereas NBS1-ATP bound conditions lost R905 contact entirely (Fig. 7b). Consistent with these contacts, TAX-R remained lower in the cavity in NBS1-ADP states and shifted toward the TM1/11 side near the P-site in ADP/ATP (Fig. 2b,c). TAX-R IF-af and TAX-H IF-mdl/IF-af showed no marked nucleotide-dependent ICL shifts, whereas TAX-P IF-mdl/IF-af showed shifts similar to TAX-R IF-mdl. For TAX-P, NBS1-ADP states showed consistent adenine–ICL contacts (R905 0.80 in ADP/ADP; D164 0.67 in ADP/ATP), whereas NBS1-ATP states showed lower ICL contact (<0.50 for all ICL residues). Similar pattern was observed in TAX-P site IF-af (Fig. 7c), where NBS1-ADP bound states showed even more persistent ICL contacts (R905 0.95 in ADP/ADP; D164 0.77, R905 1.0 in ADP/ATP) and NBS1-ATP bound conditions had moderate to low ICL contact (<0.55). In TAX-P IF-af, ICL1 W162 formed π-stacking with the adenine ring (0.37), a feature specific to this condition. These P-site contacts suggested that paclitaxel could enhance electrostatic and π-stacking interactions between ICL1/4 and NBS1-ADP. Preferential coordination of NBS1-ADP over NBS1-ATP under P-site contact supported a modulatory role for this subpocket [23]. At NBS2, R262 (ICL2) and D805/D806 (ICL3) also formed salt-bridge contacts with the adenine moiety, paralleling NBS1. ICL3 V801, F804, and P807 added hydrophobic contacts, contributing to the ICL2/3–NBS2 adenine interaction network. Although nucleotide-dependent shifts were limited, some conditions favored ICL–adenine contacts. TAX-H site IF-mdl in ADP/ADP condition displayed moderate contacts with both ICL2 (R262 0.60) and ICL3 (V801 0.53, F804 0.64, D805 0.25), which was similar in IF-af (R262 0.47, V801 0.13, D805 0.48). Notably, TAX-P IF-mdl showed low-to-moderate ICL2/3–adenine contacts (R262, V801, F804, D805; 0.10–0.48) in all states except ATP/ATP. TAX-P site IF-af displayed a similar pattern of ICL2/3 contact with nucleotide in all nucleotide states other than ATP/ATP. In IF-af, ADP/ADP and ADP/ATP showed higher contacts focused on ICL2 R262 (0.48, 0.85), and ATP/ADP engaged multiple ICL2/3 residues, R262 (0.57), V801 (0.47), F804 (0.76), D805 (0.23), D806 (0.13), P807 (0.55). Analogous to TAX-P IF-af in ADP/ATP (ICL1 W162 π-stacking with NBS1-ADP), TAX-P IF-af in ATP/ADP showed F804 π-stacking with NBS2-ADP (0.31). These ICL aromatic π-stacking interactions were specific to TAX-P IF-af and stabilized NBS1-ADP in ADP/ATP and NBS2-ADP in ATP/ADP.

A-loop aromatic residues at the NBS also played a key role in nucleotide engagement in the paclitaxel-bound systems. In NBS1, the A-loop tyrosine Y401 engaged the nucleotide adenine: explicit ⇡-stacking (Fig. 7b) reached up to 0.49, while hydrophobic contact with adenine ring reached 0.80–0.93 across conditions. Y1044 in NBS2 also engaged the adenine base: π-stacking (Fig. 7b) peaked 0.36 while adenine-ring hydrophobic contacts reached 0.56–0.63. These observations indicated a somewhat asymmetrical base recognition: the adenine moiety was more persistently stabilized by Y401 in NBS1, whereas aromatic Y1044 in NBS2 engaged the base intermittently. This aligned with the 2D contact maps, where NBS2 showed lower adenine-side contact frequency. NBS2 bound nucleotides overall displayed more disperse contact network as the adenine moiety displaced from A-loop made additional hydrophobic contacts with residues further away from the catalytic site in NBD, including V1080, F1086, Y1087, I1115, I1121. For TAX-P IF-mdl in ADP/ADP, NBS2-ADP showed moderate hydrophobic contacts with F1086 and Y1087 (0.46, 0.44). For TAX-H IF-af in ATP/ADP, NBS2-ADP exhibited low-frequency ⇡-stacking with F1086 and Y1087 (0.10, 0.09; Fig. 7b–c).

Together, these findings illustrated that both polar and non-polar interactions collectively secured the nucleotide in paclitaxel bound systems: the phosphate groups were anchored by cationic residues, while the adenine base was sandwiched by aromatic and hydrophobic side chains from the NBS and ICL regions.

### Verapamil bound IF P-gp: NBS residue–nucleotide interaction profiles

In verapamil-bound inward-facing P-gp, the nucleotide–protein interaction patterns partially mirrored those observed with paclitaxel, with some noteworthy distinctions (Fig. 8). Adenine-side contacts for NBS2-bound nucleotides were generally lower than for NBS1, though less consistently so than with paclitaxel. For example, in IF-af, NBS1-bound nucleotides showed reduced coordination overall, a pattern not seen with paclitaxel. Thus, NBS2 remained more loosely coordinated overall, whereas NBS1 coordination varied with linker conformation, implicating linker-dependent modulation. Despite linker-dependent variability at the adenine–ribose end (NBS1), phosphate contacts persisted at both NBS for most ATP/ADP states, as in paclitaxel. As with paclitaxel, a minority of states showed weaker phosphate coordination, which in verapamil runs was confined to NBS2-bound nucleotides. VER-H site IF-mdl in the ATP/ATP condition had NBS2-ATP γ-phosphate group showing inconsistent contact with NBS and VER-P site IF-mdl in ATP/ADP condition displayed NBS2-ADP α-phosphate group with lower contact frequencies. In IF-af conformers, VER-H site in ADP/ADP and ATP/ADP conditions led to lower contact frequencies for the NBS2 bound ADP α and β-phosphate groups. As with paclitaxel, phosphate-contact destabilization arose when verapamil engaged non-canonical cavity regions. For example, VER-H in ADP/ADP localized near the TM10/12 entry (Fig. 4b), and VER-P in ATP/ADP stabilized against the TM2/11 side of the P-site (Fig. S3.4i). An exception occurred for VER-H in ATP/ADP, where the contact cluster centered in the cavity (Fig. 4d) and NBS2-ATP displayed destabilized γ-phosphate contact in IF-mdl (Fig. 8b). These atom-level data supported that non-canonical drug contacts near H/P sites could weaken phosphate coordination of bound nucleotides.

**Figure 8:**
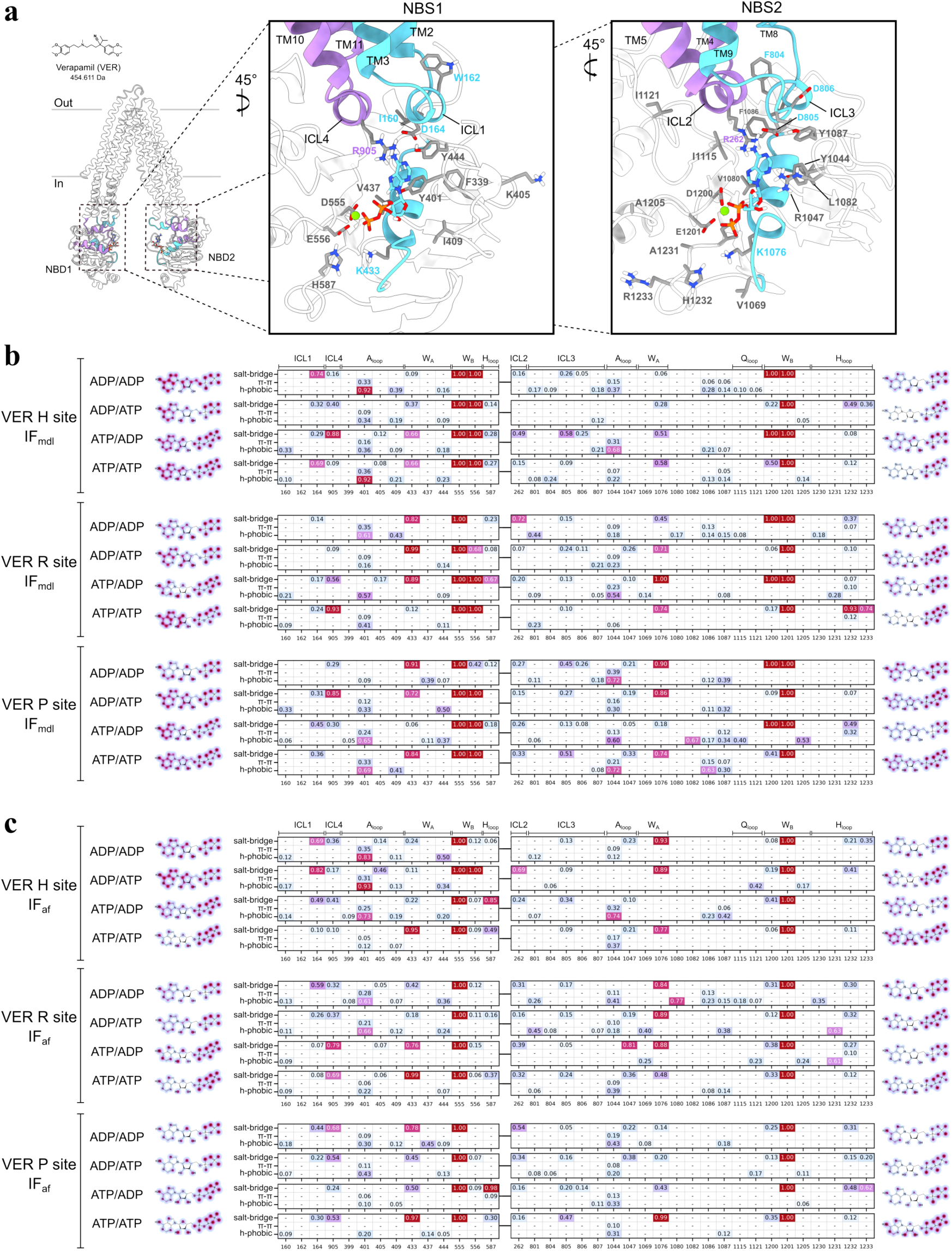
Verapamil-bound IF P-gp alters NBS–nucleotide coordination. Interaction profiles between bound nucleotides and NBS residues were calculated. (a) NBS1 and NBS2 residues are labeled; ICL1/3 and Walker A are cyan; ICL2/4 are purple. (b) Heatmaps for VER IF-mdl show salt-bridge, ⇡-stacking, and hydrophobic interaction frequencies (rows = interaction types; columns = residues), normalized over replicas. Blocks correspond to VER-H, VER-R, and VER-P IF-mdl, each with ADP/ADP, ADP/ATP, ATP/ADP, and ATP/ATP conditions. Adjacent 2D drawings contour nucleotide atoms by interaction frequency. (c) Same format for VER IF-af.

**Figure 9:**
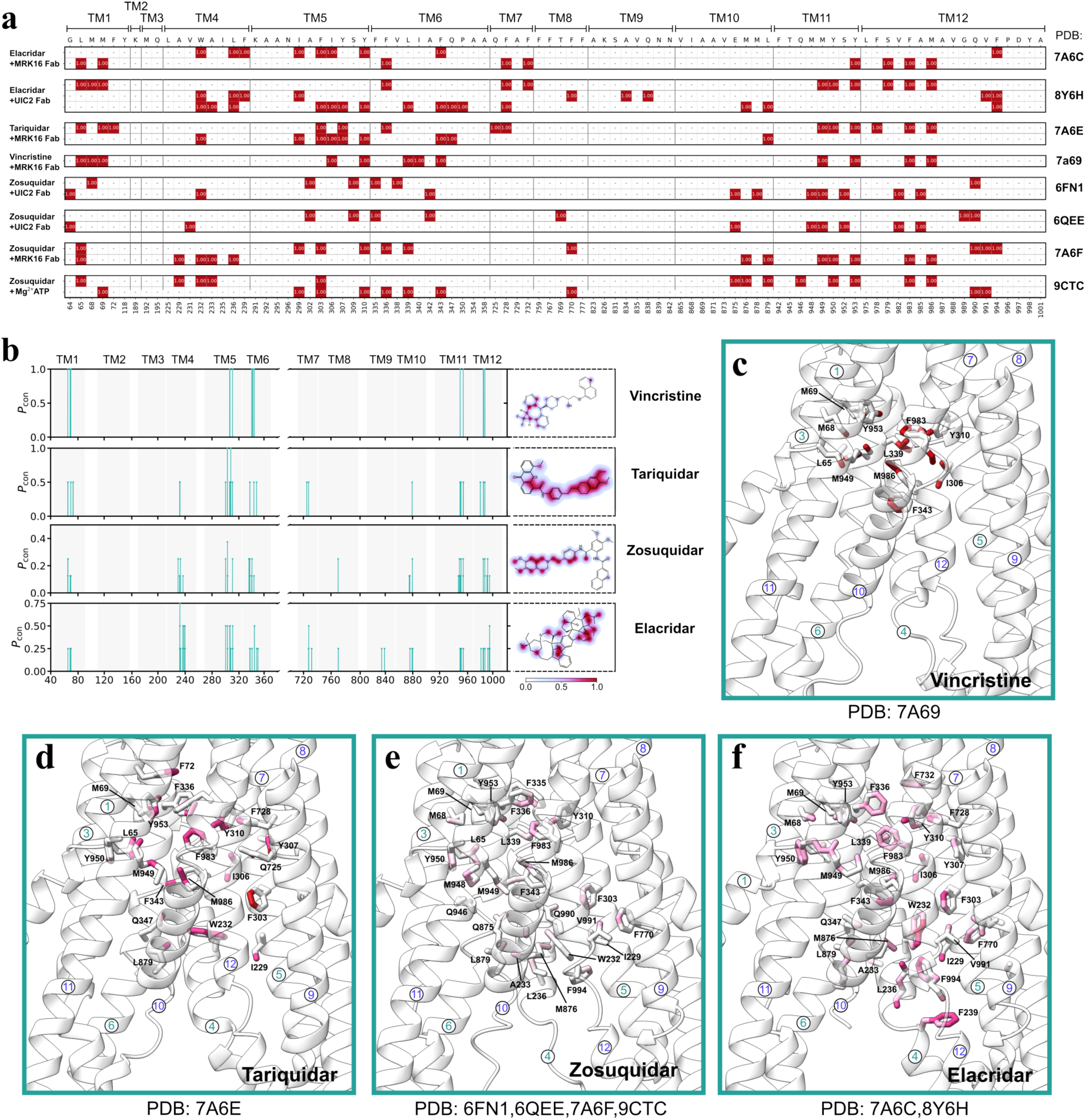
Drug-TMD interaction profile in IF P-gp calculated from previously published cryo-EM models. (a) Residue level interaction frequency between the P-gp TMD residues and the bound drug. Previously published cryo-EM structures with elacridar, tariquidar, vincristine, or zosuquidar bound were used to calculate the interactions with the protein residues. (ref) In the case of multiple molecules resolved in the cryo-EM model, the interaction pattern for each molecule was depicted separately. (b) Frequency of atom level interactions between the IF P-gp and the bound drugs colored in cyan. The ligand atoms involved in the interaction were colored with the respective interaction frequencies. (c-f) Key residues involved in the interaction with the bound drug molecule(s) as shown in panel b Residues with at least one atom with interaction frequency above 0.25 were shown.

The same cast of NBS residues was involved in coordinating the nucleotides (Fig. 8a), and the predominant interaction types remained consistent in verapamil bound P-gp compared to paclitaxel data. The Walker A lysines at NBS1 and NBS2 frequently formed high-occupancy salt-bridges to phosphates, including near-unity cases (K433 up to 0.99; K1076 up to 1.00), with lower values in other conditions. A key difference to paclitaxel bound P-gp, however, was seen with the Walker B aspartates and glutamates. In paclitaxel bound P-gp, E556/E1201 contacted the nucleotide phosphate group 100% of all frames while D555/D1200 contact frequencies varied. In verapamil bound P-gp, there was a clear nucleotide coordination asymmetry, where D555 in NBS1 and E1201 in NBS2 showed 100% contact frequency with the nucleotide phosphate groups whereas the E556 in NBS1 and D1200 in NBS2 showed variable contact frequencies. Notably, E556/D1200 contacts were reduced across IF-af simulations irrespective of cavity contacts, a pattern not seen with paclitaxel. This likely reflected different cavity-contact profiles: paclitaxel engaged larger, more confined H/R/P subpockets (Fig. 1a), whereas verapamil converged to fewer TM contacts across starts (Fig. 3a). Accordingly, more labile, fewer-residue contacts (verapamil) associated with looser nucleotide coordination at both NBS in a linker-sensitive manner. If verapamil increased NBS responsiveness to linker state, a short α-helix (residues 675–683) might have correlated with looser NBD coordination and facilitated ADP→ATP replacement at one or both NBS. The engagement of the H-loop histidines (H587 in NBS1; H1232 in NBS2) in verapamil bound P-gp showed similar pattern as in paclitaxel bound P-gp. H-loop histidines coordinated the phosphate groups of the nucleotide without any nucleotide or drug dependent pattern. Like in paclitaxel bound P-gp, NBS1 H-loop contact exclusively involved the catalytic histidine H587 whereas NBS2 H-loop contact residues involved the catalytic histidine H1232 with additional I1230, A1231 and R1233.

The intracellular coupling loop (ICL) frequently engaged the nucleotide adenine with high occupancies similar to paclitaxel bound P-gp. In NBS1 region, D164 (ICL1) and R905 (ICL4) demonstrated strong phosphate coordination in the verapamil-bound systems. R905, for example, achieved high salt-bridge frequencies (on the order of 0.8–0.9), comparable to the frequencies observed with paclitaxel. In particular, there was NBS1-ATP dependent R905-adenine coordination in VER-R site P-gp. In VER-P site IF-mdl, R905 contact frequency remained high with nucleotide adenine when NBS1 was bound to ATP (0.56 in ATP/ADP; 0.93 in ATP/ATP), while NBS1-ADP had low to no contact (<0.10). IF-af showed a similar pattern: NBS1-ATP contacted R905 at 0.79 (ATP/ADP) and 0.69 (ATP/ATP), whereas NBS1-ADP contacts were moderate (0.32–0.37). At NBS2, ICL2/3 engaged the adenine via charged and hydrophobic residues, R262 (ICL2) and V801, F804, D805, D806, P807 (ICL3), with varied frequencies. Similar to paclitaxel bound P-gp, there was no notable contact pattern that was nucleotide condition or drug-cavity contact dependent. A key difference from paclitaxel was the absence of ICL-mediated ⇡-stacking. Where P-site paclitaxel enabled ⇡-stacking by ICL1 W162 (NBS1-ADP) and ICL3/4 F804 (NBS2-ADP), verapamil showed no ICL π-stacking, consistent with a P-site contact network favored by bulkier ligands. This may have helped explain polyspecificity: bulkier, more rigid ligands could have triggered ICL ⇡-stacking with ADP and slowed turnover.

In A-loop from verapamil bound P-gp, the tyrosines were similarly involved coordinating the bound nucleotide across most nucleotide states. Unlike paclitaxel bound P-gp, A-loop coordination asymmetry between the two NBS was not observed in verapamil bound P-gp. Y401 (NBS1) engaged the adenine base: explicit ⇡-stacking (Fig. 8b) peaked 0.36, while adenine-ring contacts reached 0.83–0.89, similar to paclitaxel. Y1044 (NBS2) showed similar level of explicit ⇡-stacking frequency (peak 0.39) with adenine-ring contacts reaching 0.70–0.73. Despite reduced adenine–ribose coordination at NBS1 in some states, verapamil runs did not recruit additional NBD1 residues, unlike the broader NBD2 contacts. This suggested a more stable, less mobile NBS arrangement in NBD1 than in NBD2. NBS2 in verapamil bound P-gp still displayed additional NBD residues contacting bound nucleotides similar to paclitaxel bound systems. For instance, like in paclitaxel bound P-gp, F1086 and Y1087 in NBD2 showed low to moderate hydrophobic contacts with NBS2 nucleotide (Fig. 8), as the aromatic side chains es-tablished ⇡-stacking interaction with adenine ring. For example, VER-P site IF-mdl in ATP/ATP condition showed contact frequencies 0.15 and 0.07 between NBS2-ATP and F1086 and Y1087 respectively.

Across conformers, verapamil effects on nucleotide coordination were consistent, and key NBS interactions persisted despite IF structural nuances, similar to paclitaxel. Differences remained: NBS1 coordination and Walker B D/E involvement varied with nucleotide state and depended on linker conformation. Unlike paclitaxel, where P-site contacts increased ICL1/4 coordination of NBS1-ADP relative to NBS1, verapamil bound P-gp showed no comparable cavity-specific shifts. Instead, verapamil binding responses were linker-sensitive: IF-af showed nucleotide-dependent loosening of bound nucleotides not observed with paclitaxel. Thus, across H/R/P positions, verapamil did not impair ATP engagement at either site, maintaining symmetric nucleotide binding preference for ATP over ADP.

## 3 Discussion

It remains poorly understood how P-glycoprotein (P-gp) harnesses ATP to export a diverse array of chemically distinct substrates. Drug-free IF-wide ensembles in Chapter2 established a baseline of nucleotide-controlled heterogeneity (Fig. S2.7) that the drug-bound analysis extends with ligand-specific effects. Using two ligands with orthogonal physicochemical profiles, paclitaxel (bulky, rigid, largely hydrophobic) and verapamil (smaller, flexible, cationic), adds an additional layer of insight by contrasting packing versus electrostatics dominated recognition and their distinct consequences for NBS asymmetry and TMD–NBD coupling. Importantly, atomistic MD interrogates these effects with untethered ligands, hydrolysis-competent NBS, and a freely sampling flexible linker, thereby preserving the native allosteric communication. Key open questions include how multiple drug binding sites within the large transmembrane cavity contribute to polyspecific recognition, how ligand binding influences nucleotide coordination (and vice versa), and how structural elements like the flexible linker and intracellular coupling loops participate in the allosteric communication between transmembrane domains (TMDs) and nucleotide binding domains (NBDs). Chapter2 drug-free IF-wide simulations documented nucleotide-dependent NBS asymmetry and linker-sensitive coupling, providing a reference state for comparisons made below (Fig. 2.1c; Fig. S2.4; Fig. 2.2d) [50]. By examining paclitaxel and verapamil bound P-gp in the inward facing state, our simulations afford new insights into these issues. Comparisons to drug free IF-wide ensembles are indicated where they clarify ligand effects (Fig. 2.1c; Fig. 2.3c–d).

### 3.1 Continuous ligand cavity with nucleotide-dependent redistribution of drug binding

In our simulations, the ligand binding cavity of P-gp exhibited highly overlapping subpockets rather than rigid independent sites. Drug-free IF-wide ensembles likewise showed a contiguous cavity interior accessed by channel 1 and lateral routes (Fig. 2.3c–d), consistent with our observation of the continuous drug-bound cavity in the absence of nucleotides. We initialized paclitaxel or verapamil in three canonical binding locations (H, R, P), and during simulations the ligands sampled adjacent regions with shared TM contacts; notably, verapamil converged to the central TM6/TM12 region across the systems with differently positioned starting poses. This behavior is consistent with structural evidence that P-gp cavity can accommodate multiple ligands in partially superimposed positions [21, 59]. Paclitaxel, being a bulky and relatively rigid molecule, tended to remain closer to its initial docking pocket, whereas the smaller and flexible verapamil readily migrated toward a central cavity locus spanning both halves of the transporter. For example, paclitaxel placed in the H site persisted in a subpocket near TM4, TM5, and TM12, and with NBS2-ATP present (ADP/ATP, ATP/ATP), the drug slightly shifted upward toward the cavity center, reflected by increased contacts at central TM5 (F303/I306) and TM8 (F770) (Fig. 1a). This outcome is congruent with drug-free IF-wide simulation results, which also showed that NBS occupancy shifted interior accessibility and lateral portal prevalence, indicating that the upward movement with NBS2-ATP occurs on a background of nucleotide-shaped cavity changes (Fig. 2.3c–d). In contrast, verapamil, regardless of starting position, quickly settled into the mid-cavity, contacting a similar ensemble of residues on TM5–6 and TM11–12 across all conditions (Fig. 3a). This indicates that P-gp drug-binding region is functionally one continuous polyspecific cavity, where ligands of different shapes find partially overlapping binding modes [60, 61].

Notably, the nucleotide state modulated where each drug concentrated within the cavity. Drug-free IF-wide analyses showed analogous nucleotide-dependent shifts in A-loop distances and access pathways, indicating a shared control mechanism in the absence of ligand (Fig. 2.1c; Fig. 2.3c–d) [50]. Paclitaxel showed a nucleotide-dependent shift between an “intracellular” pose and a more “central” pose. In nucleotide-free and NBS2-ADP conditions, paclitaxel remained deep in the H-site subpocket on the TM4/5 side (e.g., TM4 F239; TM5 K291/A292/A295/N296) and engaged TM12 (F994/Y998), with W232 becoming frequent in nucleotide-bound runs; by contrast, NBS2-ATP conditions promoted more central contacts (Fig. 1a). However, when NBS1 carried ATP (ATP/ADP, ATP/ATP), R-site paclitaxel shifted toward the cavity center/upper region, reducing TM4/5 contacts and engaging upper TM6 (V/I336–I339), TM7 (F728), TM11 (Y953), and TM12 (F983) (Fig. 1a). Verapamil also redistributed with nucleotide binding, but rather than distinct distributions within different subpockets, it rearranged within the proximity of central cavity, while ligand moieties involved in cavity contact changed with translation and conformational shift of the ligand molecule. In ADP-bound states, dimethoxyphenyl groups of verapamil often nestled toward the cytosolic end of the cavity (against TM6 and TM12), whereas ATP-bound states induced the ligand to pivot upward within the cavity, for example, interacting with TM1 and TM7, while still the other side of the ligand contacting the central cavity (Fig. 3a). These motions suggest that ATP-driven conformational changes in the protein alter the cavity shape in subtle ways that favor the drug moving closer to the cavity center (and toward the eventual outward facing exit). The smaller verapamil is evidently able to dynamically reposition itself to maintain favorable contacts as the protein shifts, whereas paclitaxel, with more rigid ring systems, is prone to stay in one subpocket until a significant conformational change results in a slight positional shift. This distinction aligns with the general trend that larger, less flexible ligands (often inhibitors) form more confined high affinity poses, whereas flexible substrates can adapt within the cavity and are more readily dissociated [62, 60]. Furthermore, recent cryo EM studies show that a bulky ligand can occupy two different positions concurrently in inward facing P-gp [63, 17]. Our findings rationalize these observations: paclitaxel, like other bulky compounds, engages a subset of strong interactions in a single region (and could in principle span two adjacent pockets), while a ligand like verapamil interacts more diffusely and transiently with the cavity, making it easier to dislodge during the transport cycle.

### 3.2 Concordance with cryo-EM highlights recurrent binding hot spots

The binding patterns observed in our simulations closely mirror those seen in experimental P-gp structures with various ligands. Drug-free IF-wide ensembles overlapped IF experimental neighborhoods, which supports mapping of recurrent hot spots onto conformations present without ligand (Fig. S2.7). P-gp cryo EM complexes often show ligands bound in two main clusters within the cavity: one cluster of contacts on the TMD1 side (roughly TM4–6) and another on the TMD2 side (TM10–12), sometimes with a single ligand bridging both and in other cases with two ligand molecules each occupying one cluster [21, 63]. Our paclitaxel simulations reproducibly engaged these same clusters. For instance, at the H site paclitaxel contacted TM4 (F239 and, in nucleotide-bound runs, W232), TM5 (K291/A292/A295/N296), and, with NBS2-ATP, central TM5 residues (F303/I306), matching the two cluster pattern seen experimentally (Fig. 1a). Likewise, paclitaxel in the R-site pose contacted TM1 (L65, M69), TM6 (F343), TM11 (Y953) and TM12 (F983, M986) on the intracellular side, recapitulating the vincristine-binding pocket seen in a recent structure [17]. Across multiple ligands, the simulations recovered key aromatic and hydrophobic contacts such as F303 (TM5), F336 and F343 (TM6), Y953 (TM11) and F983/M986 (TM12) that consistently appear in cryo EM ligand complexes [64, 65]. This overlap suggests that our MD trajectories sampled the relevant binding poses corresponding to experimentally observed drug positions.

In some cases the simulations revealed additional flexibility beyond the experimental structures. For example, the cryo EM structures of verapamil-bound P-gp captured the ligand in two orientations, one primarily toward the H-site region on TMD1 and another bridging both halves in a more symmetric fashion, and our verapamil simulations encompassed both of these extremes. Moreover, we observed verapamil transiently contacting regions not prominent in the cryo EM structures, such as residues on TM7 and TM10 that lie just outside the central pocket. These extra contacts (e.g. A729 on TM7, I868 on TM10) occurred in specific nucleotide states and likely represent alternate ligand conformations that are sampled in the dynamic ensemble but may be underrepresented or averaged out in cryo EM [66]. Conversely, certain residues that appear to contact ligands in static structures were less engaged in simulation. One example is Q990 on TM12, which makes direct contact with ligands in some paclitaxel or inhibitor bound cryo EM models [26], yet paclitaxel in our trajectories rarely remained close to Q990 for long. Instead, paclitaxel favored interacting with the neighboring TM12 residues M986 and F983 that form a deeper hydrophobic pocket. This kind of discrepancy hints at the possibility of how the ligand can shift within the pocket over time, which can be further explored within a longer timescale simulation study. Overall, however, the agreement between the dominant simulation contacts and the cryo EM binding sites was strong: the major drug-binding regions identified experimentally, an aromatic-rich pocket in TMD1 and a counterpart in TMD2, clearly emerged as recurrent interaction hubs in the simulations. This convergence reinforces the idea that P-gp uses a set of nonspecific binding “hotspots” to recognize many compounds, rather than a unique site for each substrate. Polyspecificity is achieved by a ligand being able to engage one or more of these hotspots in different combinations, an interpretation supported by both mutational and structural studies [65].

### 3.3 Drug binding modulates nucleotide coordination via ICL networks with NBS asymmetry

The simulations also illuminate how drug binding influences NBD–nucleotide interactions. Drug-free IF-wide ensembles provided a baseline of NBS asymmetry against which ligand-induced changes can be evaluated (Fig. S2.3). Across conditions, the conserved motifs typically coordinated bound nucleotides: Walker A lysines (K433, K1076) and Walker B glutamates (E556, E1201) frequently formed phosphate contacts, and the A-loop tyrosines (Y401, Y1044) often engaged the adenine ring, with drug- and ligand-dependent variation. Drug-free IF-wide ensembles showed γ-phosphate-dependent strengthening of these interactions without ligand, consistent with the state-dependent variation noted here (Fig. S2.3; Fig. 2.1e). However, we observed subtle asymmetries, most clearly with paclitaxel. Its adenine ring sometimes slipped away from Y1044 and formed only transient hydrophobic contacts with surrounding residues (such as F1086/Y1087), whereas the NBS1 adenine remained sandwiched by Y401 and other contacts almost continuously. This finding aligns with biochemical data showing that the two nucleotide sites of P-gp are nonequivalent, one site typically retains ADP more weakly and exchanges nucleotides faster [67, 47]. Our simulations, especially with paclitaxel, suggest that in the inward-facing ensemble, NBS2 behaves as the more exchangeable site, with the adenine base exhibiting greater mobility when a drug is bound. Drug-free IF-wide analyses indicated tighter NBS2 geometry at IF, so greater base mobility under ligand indicates a shift relative to the drug-free reference (Fig. 2.1c).

Interestingly, specific drug–TMD interactions correlated with changes in how the intracellular coupling loops (ICLs) and NBD motifs engage the nucleotides. Drug-free IF-wide ensembles showed γ-phosphate-dependent electrostatic bridging that was NBS1 specific, as R905-nucleotide interaction appeared in high frequency when bound to ATP (Fig. S2.3). When paclitaxel occupied the P-site subpocket (near TM10/11/12), the drug-free simulation trend is reversed, as we found that ADP bound at NBS1 became more stabilized by ICL1 and ICL4 as ligand is bound on the same side of the transporter (P site located on the same side as NBS1). The P-site paclitaxel bound P-gp displayed frequent ICL–adenine contacts at NBS1-ADP showing -stacking interactions involving W162 (Fig. 7b,c), contrasted with the absence of ICL -stacking under verapamil bound conditions (Fig. 8b,c). Independent support for such bidirectional NBD–TMD communication comes from a recent study of the typeIV ABC transporter BmrA, which identified a conserved, Trp-centered “communication hinge” at the NBD–TMD interface that relays signals via intracellular coupling helix [68]. The same study showed that hinge mutations rewired local and global dynamics and could uncouple ATP hydrolysis from transport [68]. Mechanistically, this hinge-through-ICL relay aligns with our results with P-site paclitaxel bound P-gp simulations and helps rationalize the emergent NBS asymmetry. The unique NBS1-ADP stabilization feature emerging under P-site bound paclitaxel P-gp indicates that the specific contacts in the P-site allowed by the chemical properties of paclitaxel may allosterically favor the retention of ADP in NBS1 by strengthening ICL–nucleotide contacts. This mechanism is reminiscent of the “allosteric modulation” observed for certain transport inhibitors or stimulators that bias P-gp toward a state with nucleotide trapped [23, 9]. In contrast, when verapamil was bound, we did not observe the same ICL–ADP stabilization at NBS1; instead, verapamil binding led to a pattern in which NBS1-ATP remained effectively engaged. For example, R905 stayed associated with the ATP-bound NBS1 in verapamil simulation, whereas it disengaged in the equivalent paclitaxel simulation. These differences indicate that distinct drugs can differentially tune the cross-talk between the drug-binding cavity and the NBDs.

Beyond the ICLs, other NBD contacts were subtly affected by ligand binding. For example, the H-loop histidines (H587, H1232), which coordinate the γ-phosphate of ATP during catalysis, remained engaged across conditions without a robust drug-dependent pattern in our IF simulations. Drug-free IF-wide ensembles also emphasized γ-phosphate-driven coordination in IF, which aligns with this observation (Fig. S2.3). Drug binding also influenced the Walker B aspartate interaction, in paclitaxel bound trajectories the Walker B D555 at NBS1 often alternated in and out of contact with the nucleotide, whereas in verapamil-bound trajectories D555 remained fully engaged and instead its partner E556 loosened. Drug-free IF-wide ensembles did not show contacts with either of the Walker B aspartate/glutamates in both NBS (Fig. S2.3), indicating that the Walker B coordination and its divergence arises under ligand influence. Although these detailed differences require cautious interpretation, they underscore that the presence of a drug in the TMD can subtly reshape the NBS interface. This is supported by experimental evidence that drugs alter nucleotide binding and hydrolysis kinetics in P-gp [45, 46]. For instance, verapamil has been shown to increase ATP affinity at the NBDs and at high concentrations to inhibit the catalytic transition [45], while different transported substrates either stimulate or inhibit hydrolysis to varying degrees. Our simulations provide a structural context: depending on position, a ligand can stabilize or loosen overall NBS coordination, particularly the nucleotide–ICL interactions, potentially modulating turnover. Drug-free IF-wide ensembles lacked such ICL-dependent stabilization in IF, supporting the interpretation that ligand position modulates these contacts.

### 3.4 Access tunnels persist in many states and correlate with linker helicity

The inward-facing P-gp is not a static, closed container, it exhibits dynamic openings toward both the cytosol and the inner membrane. Drug-free IF-wide ensembles showed a similar pattern of cytosolic and lateral openings in the absence of ligand (Fig. 2.3c–d). Using CAVER analysis, we identified several distinct ligand-accessible tunnels linking the central cavity to the protein exterior (Fig. 5). The cytoplasmic opening (channel 1) was the predominant path in many nucleotide-bound conditions. In nucleotide-bound simulations, channel 1 often persisted at least partially (e.g., >80% of frames with ATP present still showed a continuous gap to the cytosol), indicating that even with ATP bound P-gp remained inward facing and not yet tightly occluded. In many ATP-containing runs (e.g., R-site ensembles), channel 1 persisted at high frequency (90–96%), though an IF-mdl H-site verapamil ATP/ATP case showed no detectable tunnels, indicating an occluded conformation (Fig. 5c,d). We also detected lateral openings between transmembrane helices that connect to the lipid phase. In particular, a portal between TM4 and TM6 (channels 2a/2b) and portal between TM10 and TM12 (channel 2c) were frequently observed, though their prevalence depended on the drug and the nucleotide state. For example, with R-site paclitaxel, the TM10/12 portal (2c) remained open in all nucleotide states except apo and ADP/ATP conditions, whereas with P-site paclitaxel the lateral routes (2a/2b/2c) were variable across nucleotide states (Fig. 5c, Fig. S3.5c). Drug-free IF-wide-mdl ensembles showed enhanced 2c occurrence under NBS1-ADP-containing states (Fig. 2.3c), unlike drug-bound simulations with no clear nucleotide-dependent tunnel formation trend. Verapamil, which generally resided near the cavity center, allowed mostly the TM4/6 portals to sample more occluded conformations to varying degrees (Fig. 5d). This suggests that the bulky ligand like paclitaxel occupying various subpockets within the cavity is correlated to more frequent opening of cavity toward both sides via TM4/6 and TM10/12, while smaller and more flexible ligands like verapamil which tends to primarily occupy central cavity is correlated with less frequent opening of lateral ingress tunnels, with selective opening of TM10/12 side (VER-H site-IF-mdl in ADP/ATP) and at times with complete closure of the cavity (VER-H site-IF-mdl in ATP/ATP). These findings support a model in which substrates can enter the binding cavity directly from the inner membrane leaflet via lateral gaps [62, 69, 70]. Indeed, previous simulations have shown lipids or cholesterol molecules inserting into the P-gp cavity through the TM4/6 portal [40, 66], and cysteine mutagenesis experiments confirm that both the TM4–6 and TM10–12 interfaces are functional entry gates that must open for transport to occur [71].

The flexible linker connecting NBD1 and TMD2 also appears to play a role in modulating these openings. Drug-free IF-wide ensembles linked linker helicity to association of the two transporter halves and cavity access modulation in the absence of ligand (Fig. 2.2; Fig. 2.3c,d). In our drug-bound VER-H site IF-af model, the presence of the initial helical linker segment maintained helicity and often preserved a small ligand access channel even in conditions where the IF-mdl model became fully occluded (Fig. 5d). Drug-free IF-wide ensembles previously showed similar preservation of channel 2a/2b under helical linker states in IF-wide-af (Fig. 2.2d; Fig. 2.3d). By contrast, when the linker remained unstructured in some cases (e.g., VER-H IF-mdl ATP/ATP), no tunnels were detected. This suggests that secondary structure of the flexible linker could modulate cavity occlusion propensity in the presence of a ligand bound, with a helical linker segment associated with retained access paths in some conditions. Top-cavity engagement of drug molecule further biased the linker toward helix in IF-af. Under ADP/ATP, paclitaxel at the H versus P subpockets showed distinct top-cavity contact profiles that coincided with an N-terminal-shifted helical linker (Fig.S3.3f,l); an exception was VER-R IF-af in ADP/ADP, which also displayed high helix propensity when the ligand engaged the cavity top. These subpocket-dependent effects suggest that H- and P-site engagement tunes linker helicity and, in turn, NBD dimerization efficiency, biasing IF-to-OF isomerization. Previously, drug-free IF-wide ensembles showed partial stabilization between the two opposite ICLs with linker-mediated restraints, consistent with this view (Fig. 2.1e; Fig. S2.4). Mechanistically, a helical linker likely assists NBD approach yet disfavors full dimerization, helping to avoid non-productive closure with nonhydrolyzable nucleotides. Consistent with this notion, prior studies found that P-gp remains mostly in an inward or occluded conformation until ATP hydrolysis actually occurs [72, 17].

Taken together, these dynamic tunnel and linker observations flesh out the energy landscape of the inward-facing state. Rather than a completely closed conformation, P-gp appears to continuously fluctuate, maintaining ready access for lipid soluble substrates. This behavior aligns with the “hydrophobic vacuum cleaner” model, wherein substrates partition into the inner leaflet and are then captured by P-gp [73, 62]. The inward-facing cavity can open to the membrane to admit drugs, then partially close (occlude) upon drug binding and ATP engagement, and finally open to the extracellular side to release the substrate. Our simulations are consistent with the first two stages: accessible lateral/cytosolic paths in many states and, in specific cases (e.g., VER-H IF-mdl ATP/ATP), a substate with no detectable tunnels, analogous to an occluded intermediate (Fig. 5). Full transitions to the outward-facing conformation were beyond our timescale, but the presence of occluded IF state with closed off cavity from outside while bound to ligand in the central cavity is consistent with subsequent rearrangements for exporting the drug, hinting at peristaltic pump like mechanism. The overall picture that emerges is of P-gp as a highly pliable molecular machine: drug binding, nucleotide binding, and the flexible linker together orchestrate a series of small conformational transitions that collectively result in an efficient transport cycle. By combining these results, we gain a more integrated understanding of how polyspecific drug recognition is coupled to the ATPase activity of P-gp. The simulations show that specific drug–protein contacts can regulate nucleotide handling and vice versa, which helps explain the diverse effects substrates have on the P-gp ATPase kinetics [9, 45]. They also underscore that the promiscuity of P-gp ligand cavity is enabled by a network of overlapping binding sites rather than any single “lock and key” pocket, a concept that informs how we might predict or modulate P-gp interactions [60, 74]. In summary, our findings highlight an allosteric mechanism in which drug binding and nucleotide binding reciprocally influence the coordination of each other, and structural flexibilities such as the linker and intracellular coupling helices ensure that P-gp cycles productively through conformations without getting trapped in a dead end state. Drug-free IF-wide conclusions emphasized nucleotide-driven asymmetry and linker modulation in the absence of ligand, which frame the ligand-dependent adjustments described here. This nuanced view of drug-induced dynamics in P-gp provides a framework for interpreting a range of biochemical and structural data on multidrug transporters and may guide future approaches to modulate their activity.

## 4 Conclusion

The results provide a detailed, integrative view of the initial steps of the transport mechanism in IF P-gp, highlighting the interplay between substrate binding, NBD closure propensity, and the often-neglected flexible linker. Simulations define an inward-facing ensemble of P-gp in which drug recognition arises from a continuous central cavity whose geometry and interaction network are reorganized by nucleotide state. Nucleotide occupancy redistributed contacts within this cavity (at least partially), reweighting interactions across helices rather than enforcing transitions between discrete non-communicating sites. Polyspecific binding thus reflects a malleable interior that presents overlapping subpockets tuned by nucleotide-dependent geometry. Drug engagement propagated to the nucleotide-binding domains through coupling networks that include the intracellular loops, with clear asymmetry between the two sites. This TMD-NBD coupling provides a structural rationale for differences in ATPase stimulation and inhibition across ligands, since stabilization of ADP or loosening of catalytic contacts is expected to slow turnover whereas sustained ATP engagement is expected to favor progression through the cycle. The inward-facing state maintained ligand access to cavity through cytosolic and lateral routes under many conditions, and a subset of trajectories reached occluded conformers with no detectable tunnels. The prevalence of channel 1 and of portals between TM4/6 and TM10/12 depended on nucleotide state and ligand location, consistent with capture from the inner leaflet followed by repositioning within the cavity before closure. A short α-helical segment within the linker between NBD1 and TMD2 correlated with retention of access even when other trajectories occluded, suggesting that secondary structure within this segment modulates the tendency for cavity to close prematurely. These features outline a potential energy landscape that favors drug ingress from the lipid membrane and cytosol in many microstate conformations while preserving a route toward occlusion as ATP coordinates with NBS.

An integrated mechanism therefore links a single continuous cavity, nucleotide-dependent redistribution of ligand occupancy, and asymmetric tuning of nucleotide coordination through intracellular loop networks. Linker conformation modulates access and occlusion of the cavity and nucleotide coordination, potentially reducing the risk of nonproductive closure before catalysis. The combined effect is a transport cycle that couples entry from the inner leaflet, dynamic repositioning within the cavity, selective stabilization or exchange of nucleotides, and progression toward outward facing conformation beyond the present sampling window. This framework yields testable predictions for mutational and kinetic assays and suggests design principles for modulators that bias the energy landscape, in which ligands that anchor deeply within subpockets and stabilize ADP at NBS1 are predicted to slow turnover, whereas flexible ligands that remain centrally engaged while maintaining ATP coordination are predicted to favor transport.

## 5 Materials and methods

### 5.1 Human P-glycoprotein 3D structural modeling

Crystal structures of IF murine P-gp (PDB: 4Q9H) [30], was selected as a template for building full length human P-gp homology models. The sequence in template structure was mapped to human P-gp sequence (Uniprot ID: P08183) and missing residues were modeled using Modeller v9.23 [58]. A total of 5000 models of flexible linker (residues 630-699) were generated and a conformation with the highest Discrete Optimized Protein Energy (DOPE) was selected as starting configuration. The second starting configuration of the flexible linker was constructed by adopting the AlphaFold2 predicted structure (AF-P08183-F1-v4) [57].

### 5.2 MD simulation setup

Protein was embedded in 4:1 1-palmitoyl-2-oleoyl-sn-glycero-3-phosphocholine (POPC):cholesterol bilayer using CHARMM-GUI [75] based on the protein-membrane orientation predicted by OPM database [76]. Additional minimization was performed during system generation using CHARMM-GUI to relax the membrane lipids [75].

While a coarse-grained pre-equilibration route could further homogenize the bilayer, the present atomistic protocol yielded stable protein–membrane engagement across replicas and was sufficient for the purposes of this study. ATP-Mg^2+^ were docked by aligning nucleotide bound structure of ABCB10 (PDB: 4AYT) [77]. Simulations were performed using AMBER ff14SB force field for protein [78] and LIPID14 FF [79] for POPC-cholesterol membrane, and phosphate and magnesium parameters from Meagher *et al*. [80] and Allnér *et al*. [81]. The periodic TIP3P water box was used with Na^+^ and Cl^-^ ion concentrations of 150 mM. System was energy minimized using AMBER20 [82], applying harmonic restraints with a force constant of 1000 to 0 kcal/mol Å^2^ on the heavy atoms of protein, as described in other membrane simulation protocols [56]. The rest of the simulation protocol details are identical to the procedure described in section 2.6.2.

### 5.3 CAVER tunnel analysis

The CAVER 3.0 software [83] was used to identify the possible ligand access tunnels in the simulation trajectories with frames saved at intervals of 200 ps. Initial 30% of the frames of each of the trajectories were stripped to minimize starting conformation bias. All frames of the trajectories belonging to the unique P-gp conformer and nucleotide state were combined and aligned to an initial frame. For all CAVER analyses, tunnel calculation was performed excluding the residues 1-44, 370-710, 1013-1280 to only include the TMD region. The starting points were defined as the benzene carbons in F336 and F983 to represent the canonical ligand binding site. CAVER tunnel analysis was run using a probe radius of 2.5 Å, shell radius of 6.0 Å, and shell-depth of 4.0 Å.

**Table 1:** Total number and length of initial 10 ns multi-replica MD simulations performed with drug-bound IF P-gp. All replicas were initialized from the same starting configuration with a random seed of initial velocities.

| Drug | Paclitaxel |  | Verapamil |  |
| --- | --- | --- | --- | --- |
| Conformation | IF-wide |  | IF-wide |  |
| Linker model | mdl | af | mdl | af |
| No. of nucleotide system | 5 | 5 | 5 | 5 |
| No. of drug docked sites | 3 | 3 | 3 | 3 |
| No. of replica simulation | 120 | 120 | 120 | 120 |
| Run time (ns) | 10 | 10 | 10 | 10 |
| Total run time (μs) | 18 | 18 | 18 | 18 |

**Table 2:** Total number and length of extended multi-replica MD simulations performed with drug-bound IF P-gp. Replicas showing outlier structural deviations in any predefined P-gp subdomain (see Methods) were extended to 100 ns from the initial 10 ns production run.

| Conformation | Paclitaxel |  | Verapamil |  |
| --- | --- | --- | --- | --- |
| Conformation | IF-wide |  | IF-wide |  |
| Linker model | mdl | af | mdl | af |
| No. of nucleotide system | 5 | 5 | 5 | 5 |
| No. of drug docked sites | 3 | 3 | 3 | 3 |
| No. of replica simulation | 31 | 29 | 32 | 30 |
| Run time (ns) | 100 | 100 | 100 | 100 |
| Total run time (μs) | 46.5 | 43.5 | 48 | 45 |

