## Supplementary Figures for "Drug-bound P-glycoprotein reveals nucleotide-dependent ligand pockets and ingress channels"

### 6 Supplementary data

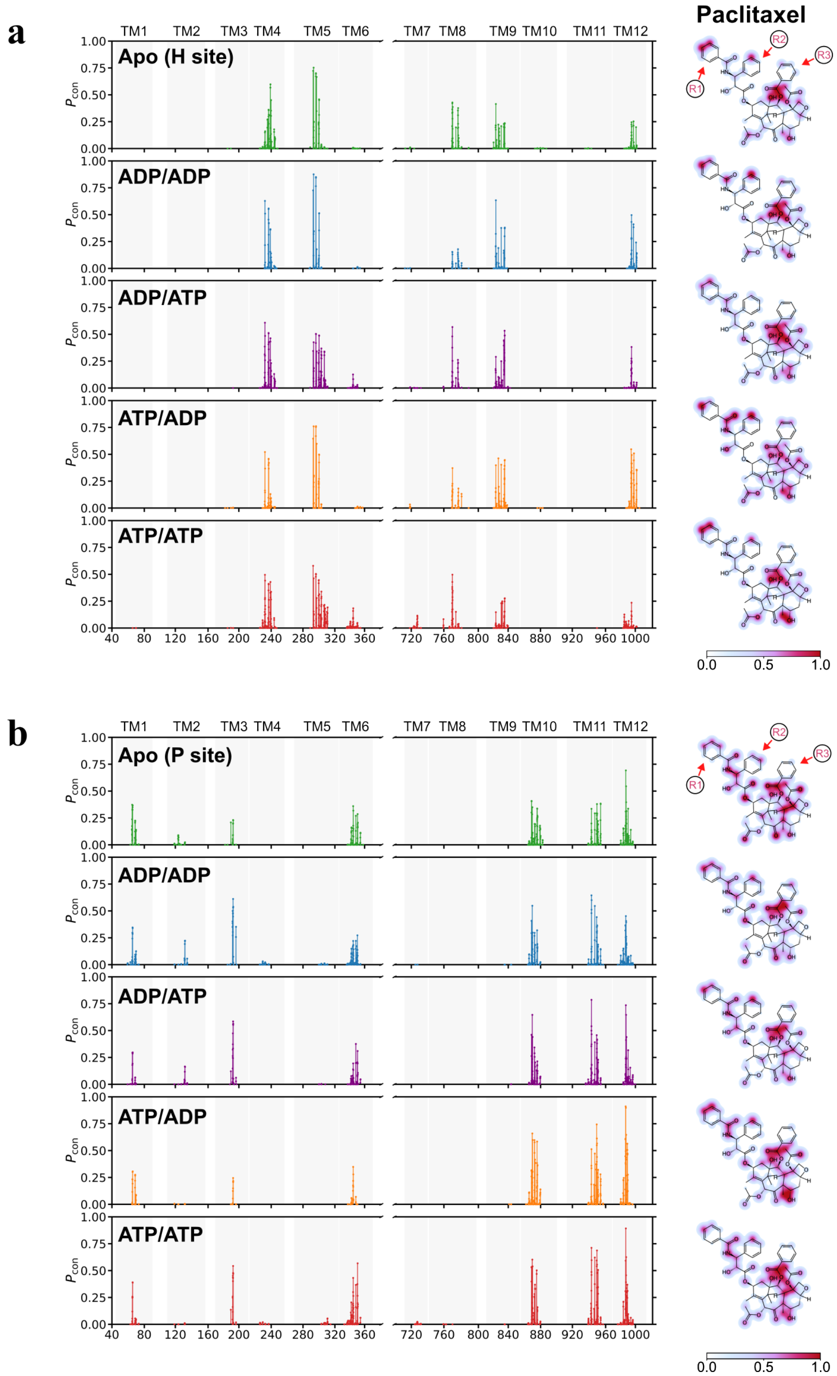


Supplementary Figure 1: Paclitaxel-TMD interaction profile in IF P-gp. (a) Frequency of atom level interactions between the IF P-gp and the bound drug paclitaxel (TAX) docked initially at the H site, with five nucleotide conditions, apo, ADP/ADP, ADP/ATP, ATP/ADP, and ATP/ATP, each colored in green, blue, purple, orange, and red, respectively. The ligand atoms involved in the interaction were colored with the respective interaction frequencies. (b) As described in panel a, but for IF P-gp initially docked with TAX at the P site.

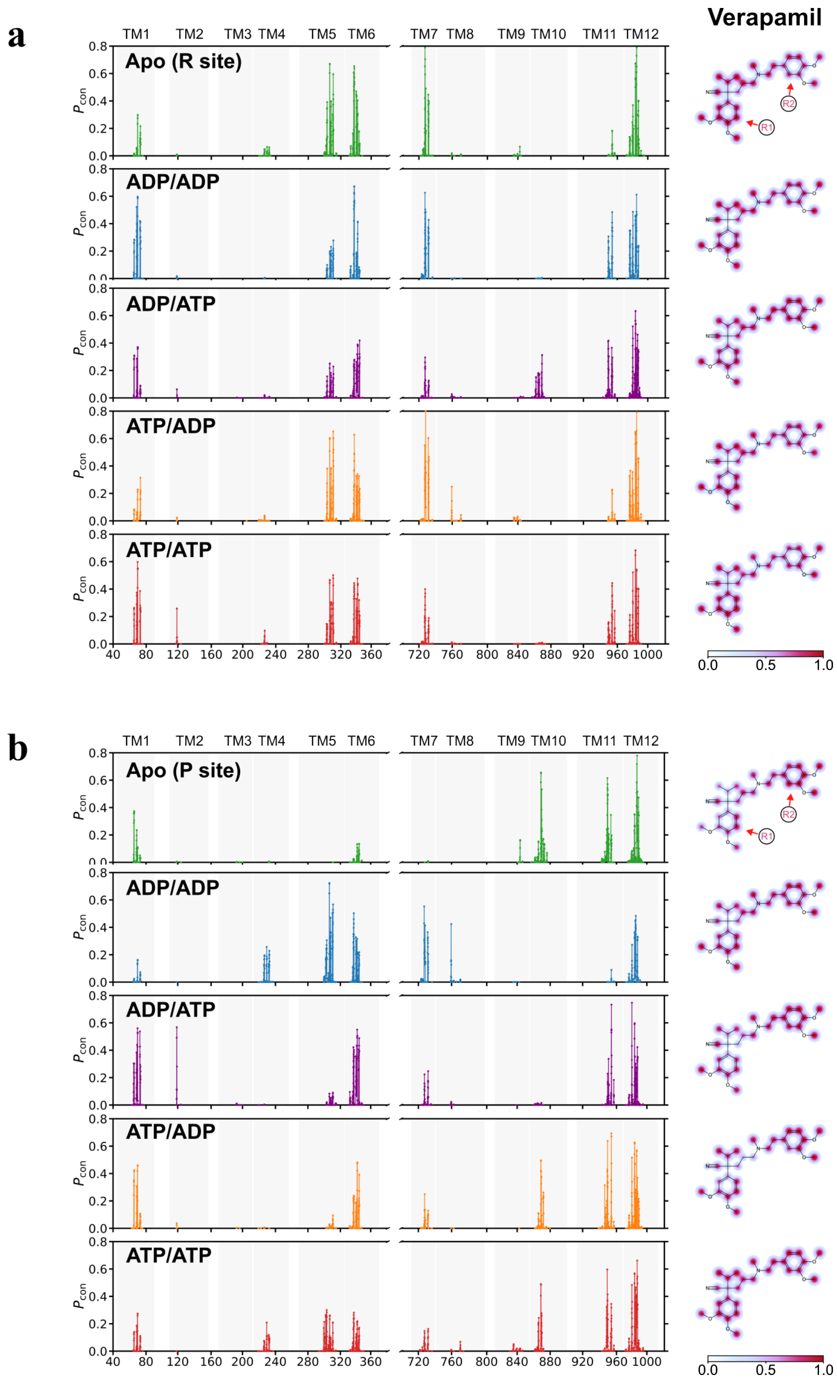


Supplementary Figure 2: Verapamil-TMD interaction profile in IF P-gp. (a) Frequency of atom level interactions between the IF P-gp and the bound drug verapamil (VER) docked initially at the R site, with five nucleotide conditions, apo, ADP/ADP, ADP/ATP, ATP/ADP, and ATP/ATP, each colored in green, blue, purple, orange, and red, respectively. The ligand atoms involved in the interaction were colored with the respective interaction frequencies. (b) As described in panel a, but for IF P-gp initially docked with VER at the P site.

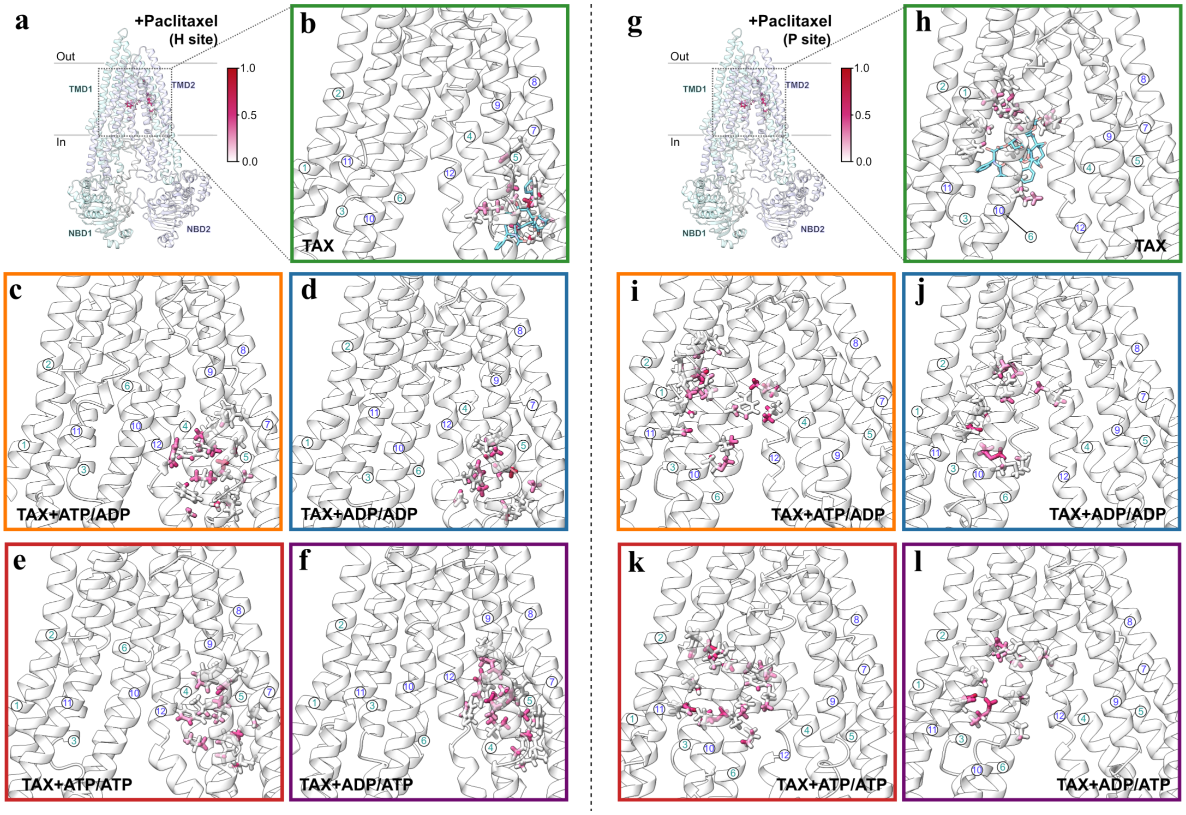


Supplementary Figure 3: Paclitaxel-TMD interaction profile in IF P-gp. (a,g) Global orientation of IF P-gp to display the area within the TMD that is analyzed in each panel. (b-f) The enlarged picture of the ligand binding site with paclitaxel (TAX) initially docked at H site shows residues involved in the interaction with verapamil in five nucleotide conditions, apo, ADP/ADP, ADP/ATP, ATP/ADP, and ATP/ATP, each in the box colored in green, blue, purple, orange, and red, respectively. Residues with at least one atom with interaction frequency above 0.25 are shown. The initial docking pose of the drug is shown in panel b. (h-l) The enlarged picture of the ligand binding site with paclitaxel (TAX) initially docked at P site. The initial docking pose of the drug is shown in panel h.

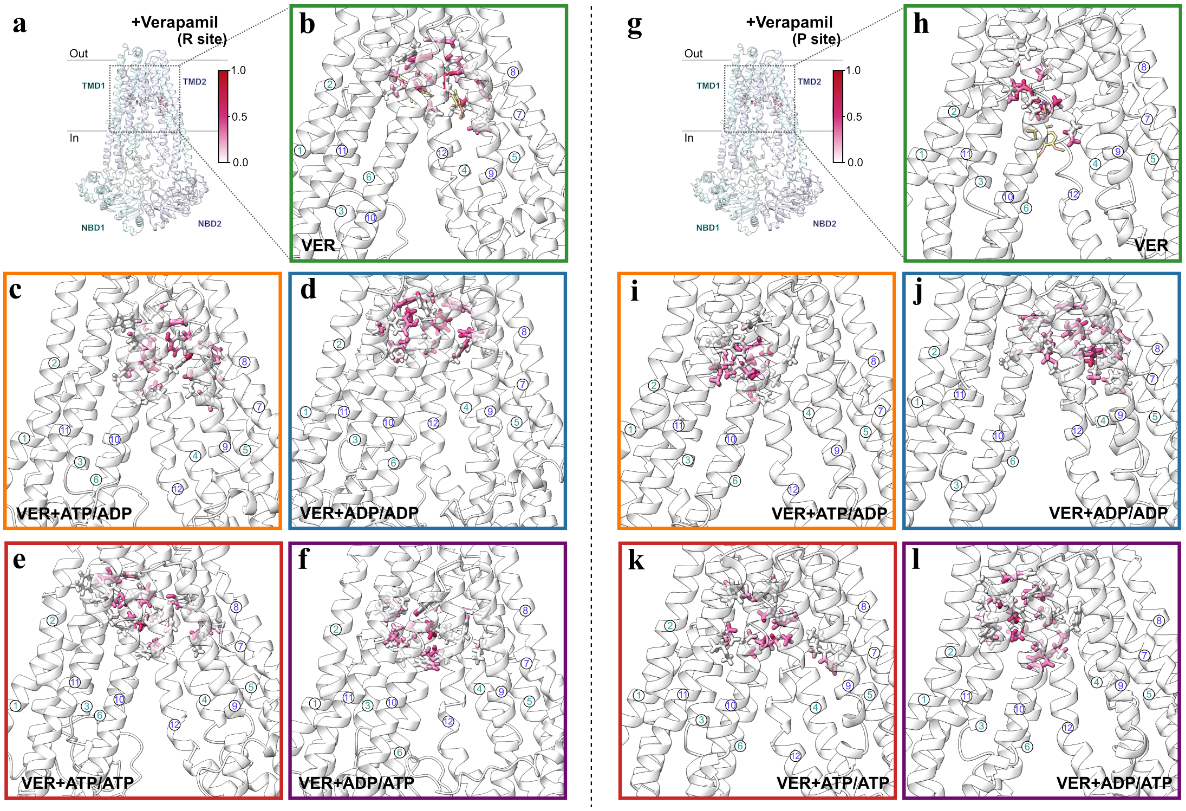


Supplementary Figure 4: Verapamil-TMD interaction profile in IF P-gp. (a,g) Global orientation of IF P-gp to display the area within the TMD that is analyzed in each panel. (b-f) The enlarged picture of the ligand binding site with verapamil (VER) initially docked at R site shows residues involved in the interaction with verapamil in five nucleotide conditions, apo, ADP/ADP, ADP/ATP, ATP/ADP, and ATP/ATP, each in the box colored in green, blue, purple, orange, and red, respectively. Residues with at least one atom with interaction frequency above 0.25 are shown. The initial docking pose of the drug is shown in panel b. (h-l) The enlarged picture of the ligand binding site with verapamil (VER) initially docked at P site. The initial docking pose of the drug is shown in panel h.

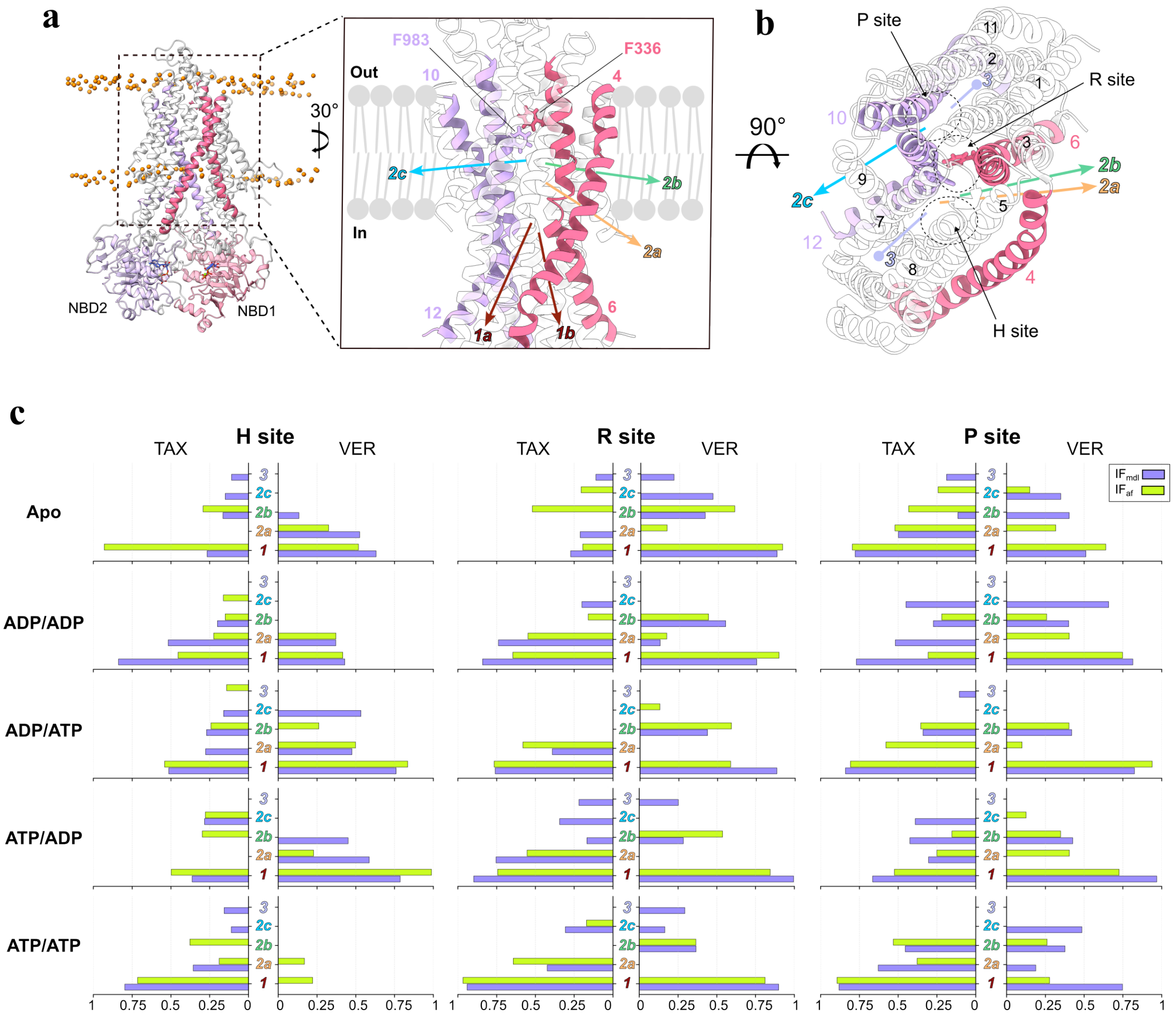


Supplementary Figure 5: Ligand-accessible tunnels computed in IF P-gp docked with paclitaxel (TAX) or verapamil (VER) at either H, R, or P site. (a) View from the lipid bilayer cross section highlighting the ligand access tunnels identified by CAVER 3.0. P-gp orientation is set to show the channel orientations in the most optimal way. NBDs may appear to be close, but are separated as theshows IF P-gp conformation. The starting positions of the probes were residues F336 and F983 within the cavity. Tunnels are defined as the following: 1a-intracellular opening near TM4/6 (brown); 1c-intracellular opening near TM10/12 (brown); 2a-intracellular side bordering lower leaflet access route via TM4/6; 2b-lower leaflet access route via TM4/6; 2c-lower leaflet access route via TM10/12; 3-inner sub-pocket near TM4/8. (b) Alternative view of the ligand access tunnels. (c) Bar plots showing the nucleotide-dependent ligand access gate formation with either TAX or VER bound IF P-gp.
